# Force-dependent activation of actin elongation factor mDia1 protects the cytoskeleton from mechanical damage and facilitates stress fiber repair

**DOI:** 10.1101/2021.02.08.430278

**Authors:** Fernando R. Valencia, Eduardo Sandoval, Jian Liu, Sergey V. Plotnikov

**Affiliations:** Department of Cell and Systems Biology, University of Toronto, Toronto, ON, Canada; Center for Cell Dynamics, Department of Cell Biology, Johns Hopkins University, Baltimore, MD

## Abstract

Plasticity of cell mechanics underlies a wide range of cell and tissue behaviors allowing cells to migrate through narrow spaces, resist shear forces, and safeguard against mechanical damage. Such plasticity depends on spatiotemporal regulation of the actomyosin cytoskeleton, but mechanisms of adaptive change in cell mechanics remain elusive. Here, we report a mechanism of mechanically activated actin polymerization at focal adhesions, specifically requiring the actin elongation factor mDia1. By combining live-cell imaging with mathematical modelling, we show that actin polymerization at focal adhesions exhibits pulsatile dynamics where spikes of mDia1 activity are triggered by contractile forces. The suppression of mDia1-mediated actin polymerization increases tension on stress fibers leading to an increased frequency of spontaneous stress fiber damage and decreased efficiency of zyxin-mediated stress fiber repair. We conclude that tension-controlled actin polymerization acts as a safety valve dampening excessive tension on the actin cytoskeleton and safeguarding stress fibers against mechanical damage.

**SUMMARY:** Valencia *et al.* report that tension-controlled actin polymerization at focal adhesions mediated by formin mDia1 controls mechanical tension on stress fibers. Suppression of mDia1 increases tension on the actin cytoskeleton leading to a higher rate of stress fiber damage and less efficient stress fiber repair.

## INTRODUCTION

All living cells are constantly exposed and respond to a myriad of dynamic mechanical cues, such as contraction, stretch, and shear, that play a central role for the control of various cellular functions including proliferation, differentiation, migration, and apoptosis (Kumar et al., 2017; Smith et al., 2017). These cellular responses are crucial to developmental morphogenesis and tissue homeostasis, as well as disease progression in cancer (Ingber, 2003; Mammoto et al., 2013; Heller and Fuchs, 2015; Michel and Dahmann, 2020). Mechanical forces generated within the cell or applied on the cell externally induce an adaptive remodeling of the actin cytoskeleton and its transmembrane linkages to the extracellular matrix (ECM), the focal adhesions, to yield changes in cell mechanics (Lecuit and Lenne, 2007; Discher et al., 2009). Internal forces produced by myosin motors pulling on actin filaments control the organization and dynamics of the cytoskeleton and focal adhesions in adherent cells (Chrzanowska-Wodnicka and Burridge, 1996; Riveline et al., 2001). External forces applied on a cell results in a strain-induced strengthening response and increase mechanical stiffness of the cell (Wang et al., 1993). This increase in cytoskeleton stiffness enhances the ability of the cells to resist mechanical injury, *e.g.* tearing of the plasma membrane and stress fibers (SFs) and strengthens cell attachment to the ECM by recruiting additional proteins to the focal adhesions, increasing the magnitude of cellular forces and the area of cell-ECM attachment (Provenzano and Keely, 2011; Murrell et al., 2015). Although the impact of mechanical forces on cell physiology is well documented, the molecular players implicated in adaptive changes in cell mechanics remain largely unknown.

Force-induced reinforcement of SFs is mostly mediated by activation of the small GTPase Rho and its downstream target Rho-associated kinase (ROCK) (Lessey et al., 2012). This signalling axis upregulates myosin light chain phosphorylation and increases cell contractility, which promotes integrin clustering and focal adhesion assembly. Upregulation of RhoA/ROCK activity also promotes actin polymerization by activating several actin nucleating factors of the formin family (*e.g.,* mDia1, FHOD1) and inhibiting the severing activity of cofilin via its phosphorylation by LIM kinase (Banerjee et al., 2020). Moreover, activation of integrins shifts the balance between capping and actin nucleating proteins in focal adhesions leading in an increase in actin polymerization and SF assembly (Vicente-Manzanares et al., 2009). Such reinforcement of the actin cytoskeleton and focal adhesions by the activity of RhoA signaling pathway is proven to be a powerful mechanism to protect cells from sustained mechanical impacts (Matthews et al., 2006). However, these responses usually occur within minutes and hours after the force is applied on the cell (Lessey et al., 2012) and thus, are unlikely to play an important role in cellular adaptation to an acute mechanical stress.

One plausible mechanism enabling cells to withstand an acute mechanical stress is by the conversion of tensile forces into a homeostatic response that rapidly remodels the SFs and dissipates excessive tension on the actin cytoskeleton. Several mechanosensitive actin-binding proteins were previously identified as common SF components indicating adaptive mechanisms finetuning the actin cytoskeleton to match the mechanical demands of cell microenvironment (Kurzawa et al., 2017). Actin polymerization at SF termini anchored at focal adhesions is essential for cellular responses to mechanical stimuli and efficient transmission of cell-generated contractile forces to the ECM (Russell et al., 2011; Tee et al., 2015; Tojkander et al., 2015; Puleo et al., 2019). In agreement with these findings, a mechanosensitive actin nucleating factor mDia1 was shown to stabilize the actin cytoskeleton and support tension through mDia1-dependent actomyosin contractility (Acharya et al., 2017; Fessenden et al., 2018), suggesting that actin nucleating factors can provide a target for cells to regulate the actin cytoskeleton upon mechanical stress.

Here, we dissected the molecular mechanism which protects the integrity of the actin cytoskeleton by dampening excessive mechanical tension on SFs. We demonstrate that actin polymerization at focal adhesions is primarily controlled by a single actin elongation factor mDia1. We show that the activity of mDia1 is modulated by contractile forces generated by the actomyosin cytoskeleton with higher forces upregulating mDia1 and increasing the rate of actin polymerization at focal adhesions. We provide evidence that the contractile forces trigger mDia1 activity resulting in pulsatile actin polymerization, which dampens mechanical tension on the SFs. Furthermore, the suppression of force-dependent actin polymerization at focal adhesions was shown to result in a two-fold increase in mechanical tension on the SFs. Such increase in tension increased the frequency of spontaneous SF damage and decreased the efficiency of zyxin-mediated SF repair. Our work reveals a novel mechanism that protects the actin cytoskeleton from mechanical damage.

## RESULTS

For multicolor live-cell microscopy of SF dynamics, mouse embryonic fibroblasts (MEFs) were co-transfected with mEmerald-β-actin and mApple-paxillin expression constructs to visualize the actin cytoskeleton and focal adhesions, respectively. In preliminary experiments, we compared the effect of either N- or C-terminally tagged β-actin overexpression on cytoskeleton organization. In agreement with previous reports (Nagasaki et al., 2017), β-actin tagged with mEmerald at the C-terminus accumulated in the cytoplasm and did not incorporate efficiently into the SFs (Fig. S1 A). Overexpression of this construct also suppressed polymerization of endogenous actin as indicated by a decrease in both the total length and number of SFs visualized by fluorescently labeled phalloidin (Fig. S1, B, and C). Conversely, moderate overexpression of β-actin tagged with mEmerald at the N-terminus had a negligible effect on the amount of total filamentous actin within the cells and the morphological parameters of the SFs (Fig. S1). Thus, N-terminally tagged β-actin was used for subsequent experiments.

### Dynamic elongation of SFs requires activity of formin family proteins

To examine the dynamics of SF elongation, we photobleached two narrow stripes across a mEmerald-β-actin labeled SF just proximal to its terminus at a focal adhesion and monitored the resultant bright spot, hereafter referred to as a photo-marker, by time-lapse spinning disk confocal microscopy (Fig. 1, A and B; Fig. S2 A). Neither width nor brightness of the photo-marker changed significantly within the 100 sec image acquisition (Fig. S2, B, C and D), however during this time the photo-marker progressively moved away from the focal adhesion at an average velocity of 0.24 ± 0.09 μm/min (Fig. 1, D, E and F and Video S1). By tracking the photo-markers and their respective focal adhesions at SF termini, we confirmed that retrograde movement of the photo-marker was not due to focal adhesion sliding (Fig. 1 C). The photo-marker movement was suppressed in cells treated with 10nM of actin polymerization inhibitor jasplakinolide, indicating that the incorporation of monomeric actin distally of the photo-marker drives SF elongation (Fig. 1, D, E and F). To assess the contribution of lateral actin incorporation in SF elongation, we measured the velocity of the photo-markers placed 2 - 10 μm away from the SF termini and found no correlation between the location of photo-marker on the SF and the rate of its movement (Fig. 1 E). Together, these data suggest that the movement of the photo-marker is mainly driven by actin polymerization at the focal adhesion resulting in SF elongation.

**Figure 1.**
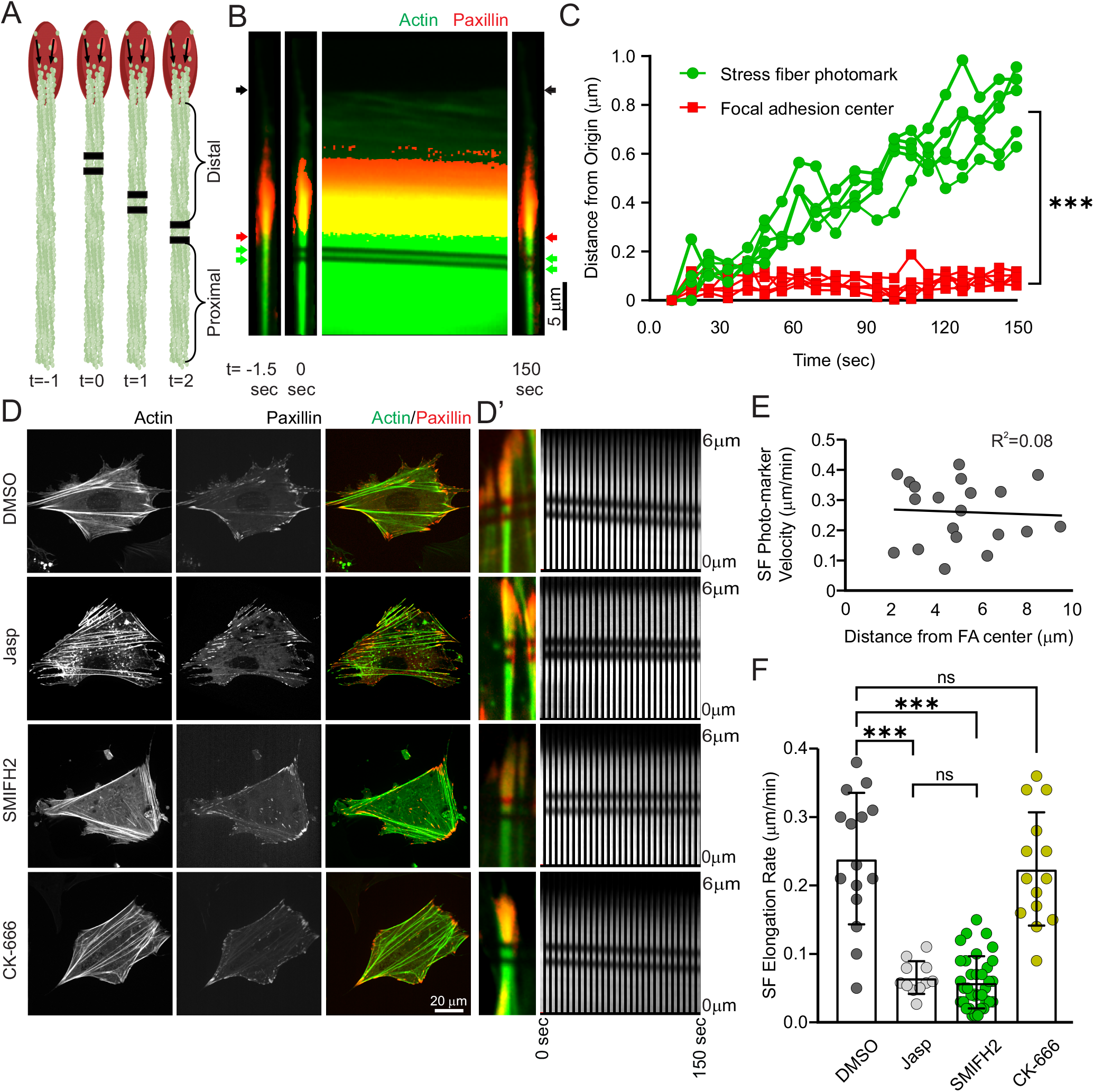
Stress fiber elongation at focal adhesions requires activity of formin family members. **(A)** Sketch of stress fiber photo-labelling by bleaching two parallel lines; actin and focal adhesion are shown in green and red, respectively. Black lines across the stress fiber demark the photo-bleached stripes. Distal and proximal SF segments relative to the photo-marker are indicated. **(B)** Close ups of an individual frames (left two panels and the most right panel) and montage (middle panel) of a stress fiber undergoing elongation at a focal adhesion. The cell was transfected with mEmerald-β-actin and mApple-paxillin 24 hrs prior to the experiment. The montage represents movement of the photo mark over 2.5 min interval. Red arrows indicate the distal tip of the focal adhesion, green arrows indicate the bleached stripes on the stress fiber, and grey arrows indicate the position of the cell edge. **(C)** Quantification of the displacement of photo-markers and the corresponding focal adhesions over 2.5 min. **(D)** Representative images of MEFs expressing mEmerald-β-actin and mApple-paxillin and treated with either 0.1% DMSO (Control), 10 nM jasplakinolide (Jasp), 10 μM SMIFH2, or 100 μM CK-666. **(D’)** Zoomed images of the photo-labelled stress fibers, and corresponding montages of the photo-markers’ movement for the experimental conditions described in (D). **(E)** Plot of photo-marker velocity *vs* distance between the fluorescent marker and focal adhesion. Grey line shows linear fit of the measurements. The regression coefficient (R^2^ = 0.08) shows no difference in stress fiber elongation rate for photo-markers located 2 – 10 μm away from the stress fiber origin. **(F)** Bar graph of the average rate of stress fiber elongation in cells treated with either 0.1% DMSO (Control), 10 nM jasplakinolide (Jasp), 10 μM SMIFH2, or 100 μM CK-666. The data on the bar graphs are presented as mean ± SD. ***, p-value < 0.001; n.s., p-value > 0.05. For C, n > 5 cells per an experimental condition. For F, n > 10 cells per an experimental condition.

To determine whether activity of actin regulators is required for SF elongation, we examined the velocity of the photo-marker movement in cells treated with either 10 μM of pan-formin inhibitor SMIFH2 or 100 μM Arp2/3 inhibitor CK-666. Consistent with previous reports (Henson et al., 2015), cells treated with 100 μM CK-666 exhibited a lack of prominent lamellipodia and a decrease in SF density in the perinuclear region accompanied with an excessive actin bunding at the cell periphery (Fig. 1 D). Cells treated for a short time with 10 μM SMIFH2 exhibited a moderate decrease in SF density compared to the controls (Figs. 1 D and S2 E). Despite the minor effect of SMIFH2 on the actin cytoskeleton, this treatment led to ≈ 75% decrease (0.06 ± 0.04 μm/min) in the rate of SF elongation at focal adhesions (Fig. 1 F). CK-666 had no noticeable effect on the rate of SF elongation (0.22 ± 0.08 μm/min), suggesting that Arp2/3 is not required for actin polymerization at focal adhesions.

Previous studies have shown that SF elongation occurs at a speed similar to that of myosin-driven actin retrograde flow (Hotulainen and Lappalainen, 2006; Endlich et al., 2007) thus we wanted to test whether suppression of SF elongation by SMIFH2 is due to a decrease in cell contractility. By Western blot analysis with a phospho-specific antibody, we examined the phosphorylation state of myosin light chain (MLC) in control and SMIFH2 treated cells. In agreement with Oakes *et al.* (Oakes et al., 2012), SMIFH2 treatment had no significant effect on the phospho-MLC level (Fig. S3 A). To test whether SMIFH2 has a local effect on SF contractility, we visualized α-actinin and phospho-MLC by immunofluorescent microscopy and analyzed the distribution of the signal along SFs. We found that α-actinin and phospho-MLC localize periodically along SFs in both control and SMIFH2 treated cells (Fig. S3 B). The intensity of phospho-MLC staining was also unaffected, indicating that the decrease in SF elongation rate in SMIFH2-treated cells is not due to a suppression of myosin contractility. Together, these data demonstrate that SF elongation at focal adhesions is driven by the activity of actin elongating factors of the formin family.

### Formin siRNA screen reveals mDia1 as a potent regulator of SF elongation

Several members of the formin family (Dia1, Dia2, FHOD1) have been previously reported to promote SF assembly by facilitating actin polymerization at focal adhesions (Hotulainen and Lappalainen, 2006; Gupton et al., 2007; Schulze et al., 2014). While it has never been shown that three other members of the family (Daam1, INF2 and FMN2) regulate SF assembly, these formins have been identified as focal adhesion components by biochemical assays and therefore are potentially implicated in SF elongation at focal adhesions (Higashi et al., 2010; Kuo et al., 2011). By using reverse transcription PCR, we first assessed the expression of the formin candidates listed above in MEF cells (data not shown). Out of six candidates, FMN2 was the only formin with an undetectable expression in MEFs, which is consistent with a prevalent expression of this protein in neuronal cells (Law et al., 2014; Sahasrabudhe et al., 2016). Other candidates (mDia1, mDia2, DAAM1, FHOD1, and INF2) were expressed in MEFs and thus, potentially facilitate actin polymerization at focal adhesions.

Previous studies have shown that actin polymerization is required for the assembly of SFs and focal adhesions in migrating cells (Kim and Wirtz, 2013). Thus, we first sought to examine how altering expression of formins that potentially facilitate actin dynamics at focal adhesions affects the architecture of SFs and maturation of focal adhesions. We transfected MEFs with siRNAs targeting mDia1, mDia2, DAAM1, FHOD1, or INF2 and co-stained the cells with fluorescently labeled phalloidin and an antibody against paxillin. The staining did not reveal any gross morphological difference in cell shape between control and knock down (KD) cells (Fig. 2, A and B). However, an unbiased morphometric analysis revealed a marked decrease (by ≈50%, p value < 0.05) in cell area, number of SFs and total length of SFs in mDia1, mDia2, and DAAM1 KDs compared to control cells (Fig. 2, C and D). Consistent with these results, the KDs conferred a significant reduction in the number of focal adhesions and total adhesion area per cell (Fig 2, A and E). While depletion of FHOD1 and INF2 led to a suppression of focal adhesion maturation as indicated by an increased fraction of nascent adhesions (Fig. 2, E and F), the morphometry of SFs in INF2 and FHOD1 KDs was similar to that of control cells (Fig. 2 D), suggesting that INF2 and FHOD1 are largely dispensable for SF assembly in MEFs and leaving mDia1, mDia2 and DAAM1 as plausible candidates to assemble SFs by facilitating actin dynamics at focal adhesions.

**Figure 2.**
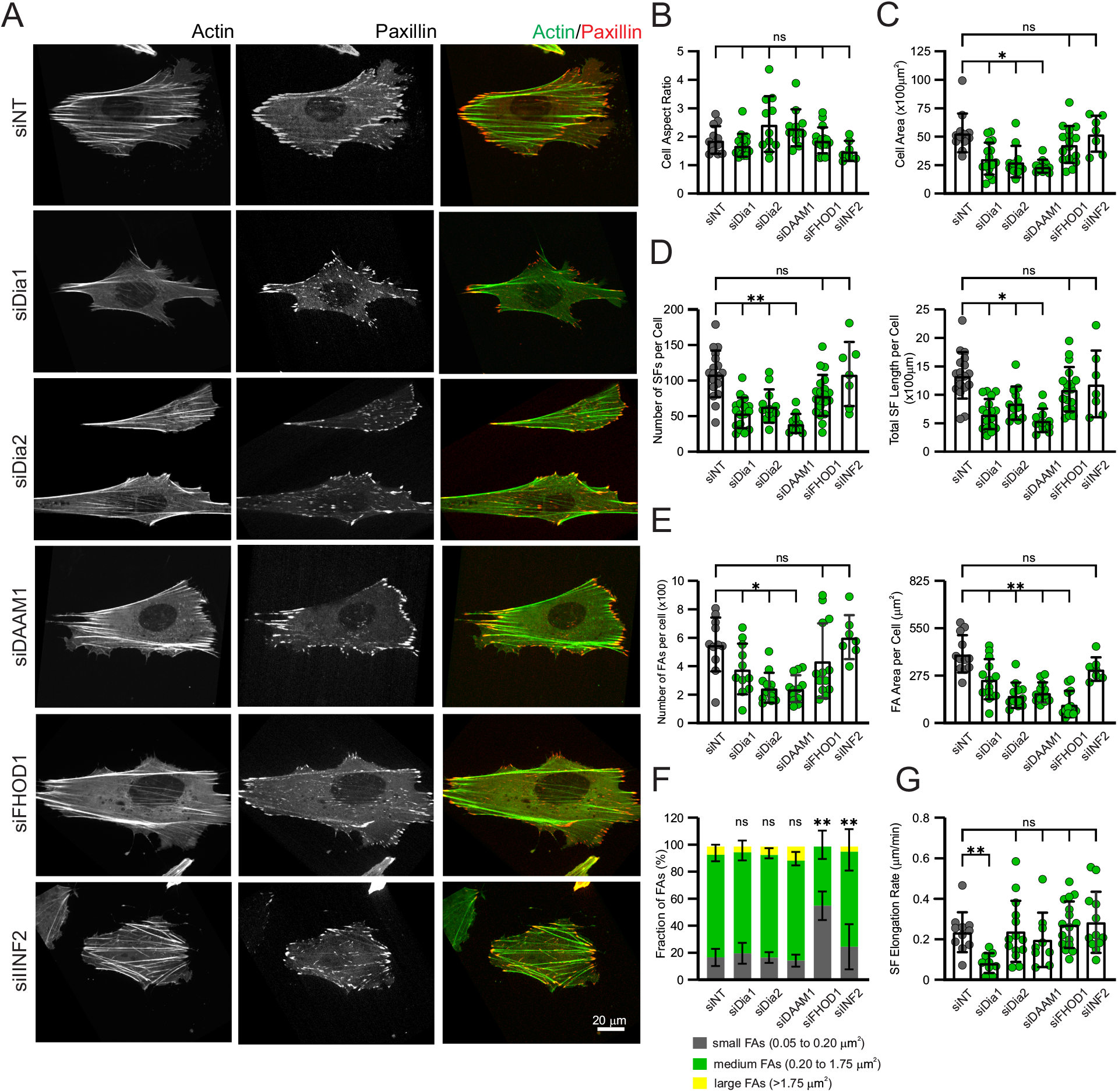
Formin siRNA screen reveals mDia1 as a potent regulator of stress fiber elongation at focal adhesions. **(A - F)** MEFs were transfected with indicated siRNAs for 24 hrs and immunostained for filamentous actin and paxillin. **(A)** Representative images of the actin cytoskeleton (left column) and focal adhesions (middle column). Images on the right show color overlay of the actin and focal adhesion channels. **(B, C)** Quantification of cell morphology upon depletion of the indicated formin family members. **(B)** Bar graph of the average cell area. **(C)** Bar graph of the cell shape factor calculated as a ratio of cell length to the cell width. **(D)** Quantification of actin architecture upon depletion of the indicated formins. The average number of stress fibers per cell is shown on the left panel. Bar graph on the right shows the total length of the stress fibers per cell. **(E, F)** Quantitative analysis of focal adhesions in the cells transfected with siRNAs targeting indicated formins. **(E)** Bar graphs of the average number of focal adhesions per cell (left panel) and the total focal adhesion area per cell (right panel). **(F)** Stacked bar graph of the percentage of small, medium, and large focal adhesions in MEFs transfected with the indicated siRNAs. **(G)** Bar graph of the average rate of stress fiber elongation in MEFs co-transfected with mEmerald-β-actin, mApple-paxillin and indicated siRNAs. The data are presented as mean ± SD. **, p-value < 0. 01; *, p-value < 0.05; n.s., p-value > 0.05. Scale bar, 20 μm. For B - F, n > 10 cells per an experimental condition. For G, n > 10 cells per an experimental condition.

To directly examine the role of mDia1, mDia2 and DAAM1 in actin dynamics at focal adhesions, we altered the expression of these formins by using siRNA and measured the rate of SF elongation by tracking the retrograde movement of the fluorescent photo-marker as described above. While depletion of mDia2 and DAAM2 had a negligible effect on the velocity of the photo-marker, cells transfected with mDia1-targeting siRNA exhibited a significant decrease (by ≈72%, p value < 0.01) in SF elongation rate compared to the controls (Fig. 2 G). The suppression of SF elongation was not due to a change in focal adhesion maturation state as indicated by a similar fraction of nascent and mature focal adhesions in control and mDia1 KD cells (Fig. 2 F). The rate of SF elongation in mDia1 depleted cells (0.07 ± 0.01 μm/min) was remarkably similar to that in the cells treated with 10 μM SMIFH2 (0.08 ± 0.01μm/min), suggesting that mDia1 is the only formin facilitating actin polymerization at focal adhesions.

### SF elongation at focal adhesions requires mDia1

To further investigate the role of mDia1 in SF elongation at focal adhesions, we first visualized mDia1 and focal adhesions by immunofluorescent microscopy. In agreement with previous reports (Carramusa et al., 2007; Higashi et al., 2019), mDia1 was uniformly distributed in the cytoplasm, but was slightly enriched at focal adhesions (Fig. 3 A). By using smart pool siRNA (see Methods for details), we decreased mDia1 protein level by ≈85% and measured the rate of SF elongation by tracking the retrograde movement of the photo-marker as described above (Figs. 3, B, C and S3 C). These experiments showed that depletion of mDia1 resulted in ≈70% decrease in SF elongation rate compared to controls transfected with non-targeting siRNA (Fig. 3 D and Video S2). Similar to SMIFH2 treatment, depletion of mDia1 led to a moderate decrease in the number of focal adhesions and SFs (Fig. S3, D and E), but likely had no significant effect on the mechanochemical properties of the SFs as indicated by similar morphology and protein composition of the SFs is control and mDia1 KD cells (Fig. S3 F and not shown). Re-expression of human Dia1 (and thus refractory to siRNA targeting the mouse protein) tagged with BFP in mDia1 KD cells fully rescued the velocity of the photo-marker and the organization of the actomyosin cytoskeleton (Figs. 3 D and S3, D, E, and F and Video S3). Together, these data show that actin polymerization mediated by mDia1 drives SF elongation at focal adhesions.

**Figure 3.**
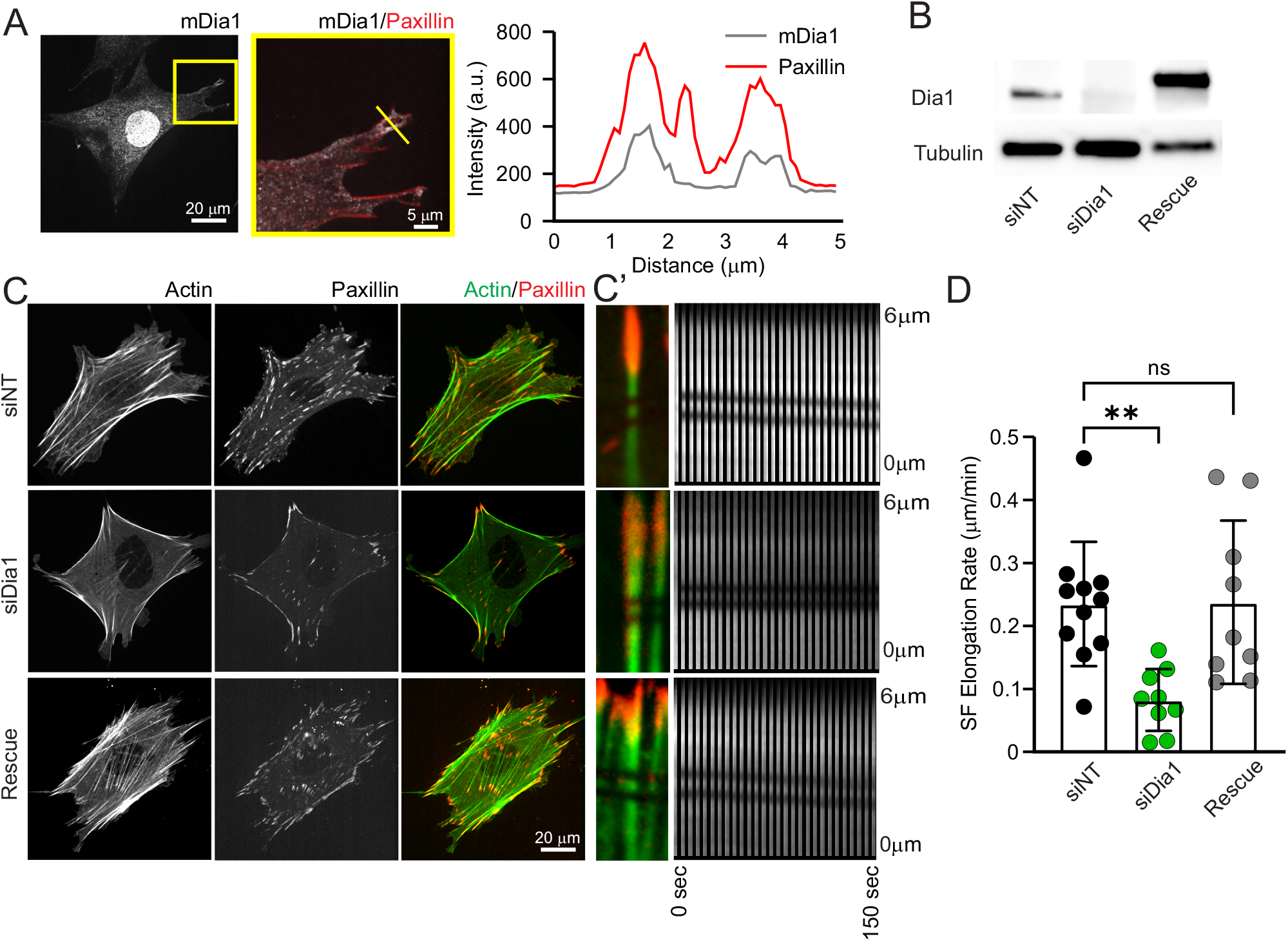
Stress fiber elongation at focal adhesion requires mDia1. **(A)** Immunofluorescent image of a MEF cell stained for mDia1 and paxillin. The yellow box on the mDia1 image indicates a region of the cell that is zoomed-in. **(A’)** Profile of fluorescence intensity for Dia1 (grey line) and paxillin (red line) channels along the yellow line. **(B)** Western blot of MEFs transfected with non-targeting siRNA (siNT) or mDia1-targeting siRNA (siDia1 and Rescue conditions). Cells shown for the rescue condition were also transfected with an expression construct for siRNA resistant human Dia1 (hDia1-BFP). Western blot was probed against αTubulin and Dia1 **(C)** Representative images of siNT, mDia1-depleted, and hDia1-rescued MEFs expressing mEmerald-β-actin (left panels) and mApple-paxillin (middle panel). **(D)** Zoomed images of photo-labelled stress fibers, and corresponding montages of photo-markers’ movement in siNT, mDia1-depleted, and hDia1-rescued MEFs. **(F)** Bar graph of the average rate of stress fiber elongation in siNT, mDia1-depleted, and hDia1-rescued MEFs. The data are presented as mean ± SD. **, p-value < 0. 01; n.s., p-value > 0.05. For D, n > 15 cells per experimental condition.

To determine whether the role of Dia1 in control of actin polymerization at focal adhesion is specific to MEFs, we measured the rate of SF elongation in primary human gingival fibroblasts, as well as in U2OS and OVCAR cancer cells upon depletion of Dia1. In agreement with previous reports (Hotulainen and Lappalainen, 2006; Endlich et al., 2007), these experiments revealed a minor difference in SF elongation rate among different cell types (Fig S3 G). However, all tested cell lines exhibited a drastic (~70%) decrease in SF elongation rate when treated with Dia1-targeting siRNA. These data along with the report from the Lappalainen group demonstrate the conserve function of Dia1 in facilitating SF elongation across various cell types.

### mDia1-dependent SF elongation requires ROCK-mediated actomyosin contractility

Previous studies have demonstrated an upregulation of mDia1 activity by mechanical tension (Jégou et al., 2013; Yu et al., 2017). Thus, we thought to examine whether mDia1-dependent SF elongation is regulated by p160 Rho kinase (ROCK)-dependent contractility. We partially inhibited the activity of ROCK by treating MEFs co-expressing mEmerald-β-actin and mApple-paxillin with a low concentration (2 μM) of Y-27632 (“ROCK inhibitor”) and measured the rate of SF elongation by tracking the retrograde movement of the photo-marker as described above. Consistent with previous reports (Katoh et al., 2011; Stricker et al., 2013), the SFs and focal adhesions assembled in cells treated with 2 μM ROCK inhibitor were not statistically significant from the ones in control cells (Fig. 4, A, B and C). However, this treatment caused a ≈70% decrease in the rate of SF elongation compared to control cells (Fig. 4 D). Cells treated with ROCK inhibitor exhibited a similar value for the SF elongation rate as for mDia1 KD cells in the absence of ROCK inhibitor. Since the activity of mDia1 does not require ROCK (Carramusa et al., 2007; Brandt et al., 2007; Kühn and Geyer, 2014), the effect of the ROCK inhibitor on SF elongation is not likely to be attributed to a direct suppression of formin activity by the inhibitor and suggests that ROCK-mediated contractility facilitates mDia1-mediated actin dynamics at focal adhesions.

**Figure 4.**
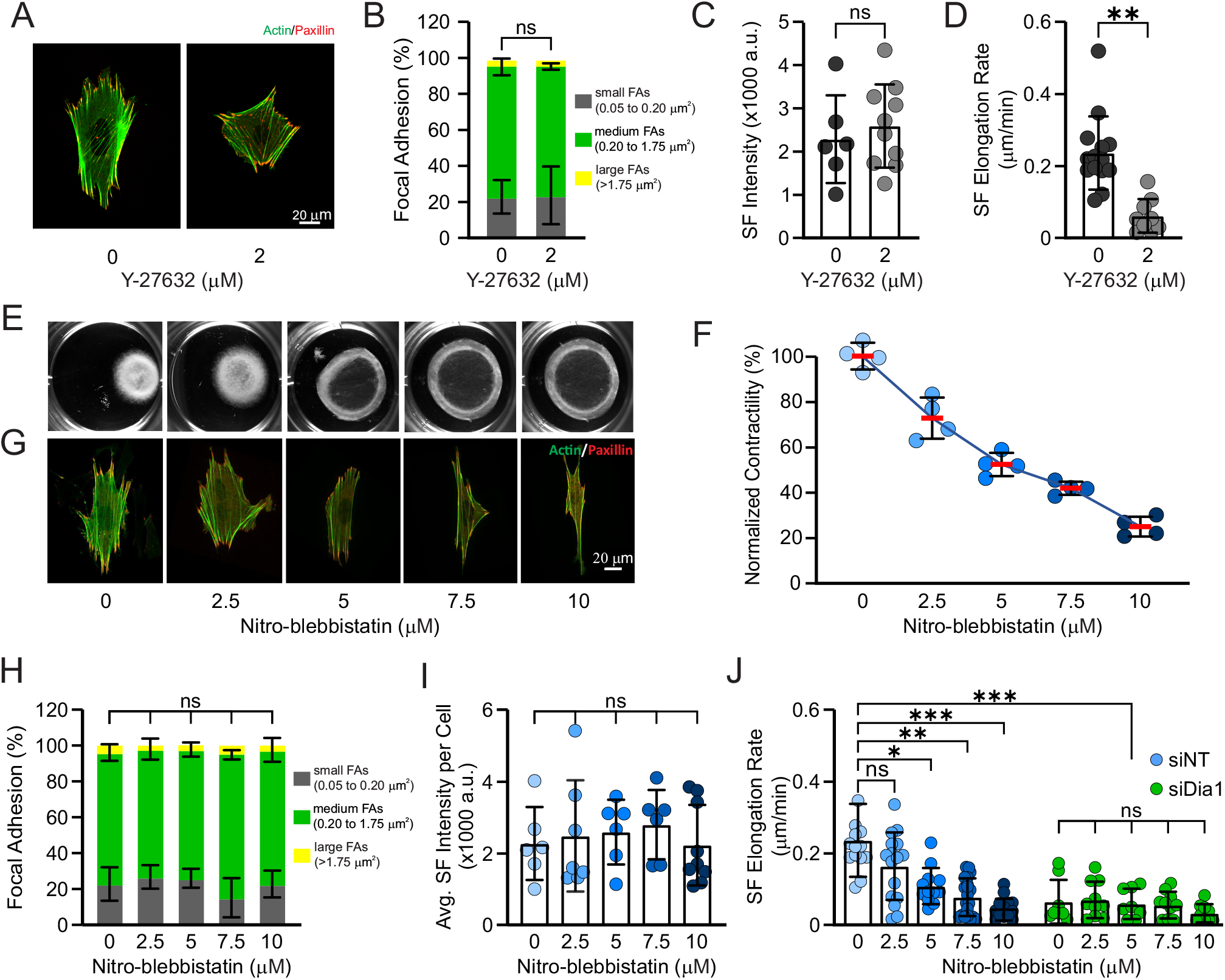
Stress fiber elongation at focal adhesions requires ROCK-mediated actomyosin contractility. **(A)** Representative images of MEFs treated with 0.1% DMSO (vehicle) or 2 μM Y-27632 for 2 hrs and immunostained for filamentous actin and paxillin. **(B, C)** Quantitative analysis of focal adhesions and stress fibers in control and ROCK-inhibited cells. **(B)** Stacked bar graph of the percentage of small, medium, and large focal adhesions. (**C)** Bar graph of the average fluorescence intensity of stress fibers stained with phalloidin-AlexaFluor488. **(D)** Bar graph of the average rate of stress fiber elongation in cells treated with 0.1% DMSO (vehicle) or 2 μM Y-27632. **(E)** Images of collagen gels contracted by MEFs in the presence of the indicated concentrations of nitro-blebbistatin. **(F)** Quantification of the collagen gel contraction induced by MEFs in the presence of 0 – 10 μM nitro-blebbistatin. Normalized contractility was calculated by dividing the area of the collagen gel in the absence of nitoblebbistatin by the area of the collagen gel in the presence of a given nitro-blebbistatin concentration (n > 4 independent assays). **(G)** Representative images of MEFs cultured on fibronectin-coated coverslips in the presence of the indicated concentration of nitro-blebbistatin and immunostained for filamentous actin and paxillin. **(H, I)** Quantitative analysis of focal adhesions and stress fibers in control and myosin-inhibited cells. **(H)** Stacked bar graph of the percentage of small, medium, and large focal adhesions. **(I)** Bar graph of the average fluorescence intensity of stress fibers stained with phalloidin-AlexaFluor488. **(J)** Bar graph of the average rate of stress fiber elongation in control (siNT, blue dots) and mDia1 depleted (siDia1, green dots) cells treated with the indicated concentration of nitro-blebbistatin. The data are presented as mean ± SD. ***, p-value < 0.001; **, p-value < 0. 01; *, p-value < 0.05; n.s., p-value > 0.05. n > 6 cells per experimental condition.

To further examine the role of actomyosin generated contractile forces in SF elongation, we used a photostable small molecule inhibitor of myosin II, (S)-nitro-blebbistatin (thereafter referred as nitro-blebbistatin). We first assessed the effect of nitro-blebbistatin on cell contractility and gauged the effective inhibitor concentrations by culturing MEFs in free-floating collagen gels in the presence of 2.5 – 10 μM nitro-blebbistatin, following by quantification of gel contraction. Increasing concentration of nitro-blebbistatin resulted in a progressive decline in cell contractility that reached ≈83% for the cells treated with 10 μM nitro-blebbistatin (Fig. 4, E, F and G). Like the effect on SFs, nitro-blebbistatin treatments caused a significant decrease in the number of focal adhesions per cell but had no effect on the average size of focal adhesions and fraction of mature focal adhesions (Figs. 4 H and S4 A). Remarkably, all tested concentrations of nitro-blebbistatin had a limited effect on organization of the SFs visualized by immunofluorescent microscopy: while causing a ≈20-50% decrease in the number of SFs per cell, these treatments had no effect on the mean intensity of phalloidin staining of the SFs (Figs. 4, I and J and S4 B). Together, these data show that partial inhibition of myosin II with nitro-blebbistatin does not prevent focal adhesion and SF assembly.

Next, we used nitro-blebbistatin treatment to determine the role of myosin contractility in the SF elongation mediated by mDia1. We treated control and mDia1 KD cells expressing mEmerald-β-actin and mApple-paxillin with a range of nitro-blebbistatin concentrations, labeled individual SFs with a fluorescent photo-marker and measured the rate of SF elongation by tracking the retrograde movement of the photo-marker as described above. This analysis revealed a dose-dependent decrease in the rate of SF elongation in control cells treated with non-targeting siRNA as the concentration of nitro-blebbistatin increased (Fig. 4 J). In contrast, the rate of SF elongation in mDia1 KD cells was insensitive to nitro-blebbistatin – SFs elongated at 0.075 ± 0.01 μm/min regardless of the presence of nitro-blebbistatin. Together these data demonstrate that contractile forces facilitate actin polymerization mediated by mDia1 at focal adhesions.

### Contractile forces trigger cyclic mDia1-dependent actin polymerization at focal adhesions

To gain mechanistic insight into how contractile forces govern the dynamics of SF elongation, we first developed a mechanical model of actin polymerization at a stationary adhesion site. The current model includes three functional modules: (*i*) a stationary focal adhesion, (*ii*) a bundle of unidirectionally polarized actin filaments anchored at the barbed ends at the focal adhesion and pulled by myosin motors at the pointed ends, and (*iii*) a force-sensitive actin polymerization module associated with the barbed ends of the actin filaments (Fig. 5, A; the model description is provided in Methods).

**Figure 5.**
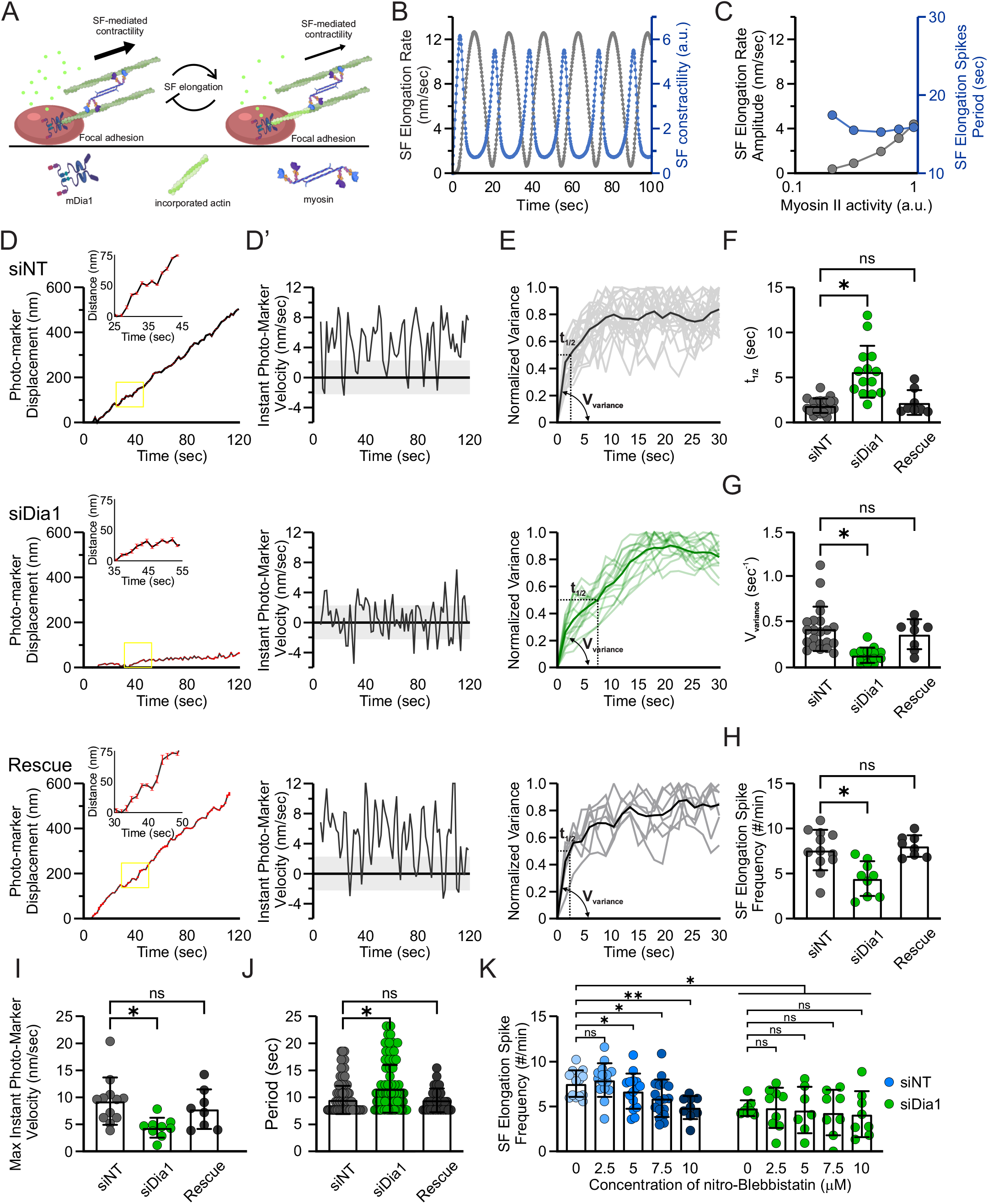
Actomyosin-generated contractile forces trigger mDia1 activity at focal adhesion and induce cycles of stress fiber elongation. **(A)** Schematic description of the mathematical model where myosin (purple) generates tension and pulls on existing stress fibers (dark green), stress fibers terminate at focal adhesion complexes (red) containing mDia1 (blue). Stress fiber-focal adhesion interface in a high-tension state is shown on the left. High tension increases the activity of actin polymerization module and facilitates incorporation of globular actin monomers to the barbed ends of actin filaments and stress fiber elongation (shown on the right). Such incorporation of actin monomers decreases tension on the stress fiber and supress actin polymerization module. **(B)** Plot of stress fiber elongation rate (grey) and actomyosin-generated contractile forces (blue) over time predicted by the mathematical model. **(C)** Plot of the maximum stress fiber elongation rate (grey) and the duration of actin polymerization cycles (blue) as a function of cellular contractility predicted by the mathematical model. **(D)** Representative traces of the photo-marker movement in cells transfected with non-targeting siRNA (siNT) or mDia1-targeting siRNA (siDia1 and Rescue). For the rescue conditions cells were additionally transfected with siRNA resistant hDia1-BFP expression construct. A yellow square represents a zoomed in region of the photo-marker trace. **(D’**) Corresponding plots of instantaneous photo-marker velocity over time for the conditions indicated in (D). Grey box indicates the range of photo-marker velocity that is not significantly different from the experimental noise. **(E)** Plot of the variance in instantaneous photo-marker velocity over time for the experimental conditions indicated in (D). Grey and pale green lines represent variances for individual photo-markers. Thick black and green lines represent the averaged variance for a given experimental condition. **(F, G)** Bar graphs of t1/2 (F) and slope (G) of the variance curves for the experimental conditions indicated in (D). **(H - J)** Quantification of the cycles of stress fiber elongation for the conditions indicated in (D). **(H)** Bar graphs of the average frequency of stress fiber elongation spikes. **(I)** Bar graph of the average maximum instantaneous velocity of the photo-marker. **(J)** Bar graph of the average duration of the spikes of stress fiber elongation. **(K)** Bar graph of the average frequency of stress fiber elongation spikes in control (siNT, blue dots) and mDia1 depleted (siDia1, green dots) cells treated with the indicated concentration of nitro-blebbistatin. The data are presented as mean ± SD. **, p-value < 0. 01; *, p-value < 0.05; n.s., p-value > 0.05. n > 6 cells per experimental condition.

The dynamic model runs from the time t_*0*_ = 0 sec to the t_*end*_ = 100 sec, with a step size t_*delta*_ = 0.1 sec. In the absence of force at time t_*0*_, the baseline activity (Z_*0*_) of actin polymerization module provides a slow actin assembly at the adhesion site by adding actin monomers to the barbed ends of actin filaments from an unlimited pool of profilin-G-actin complex and elongates the actin bundle at a rate of ≈ 1 nm/sec. As the model runs, at every time step, multiple myosin motors (N_*m*_) exert a force F_*m*_ on the actin bundle that must be balanced by tension and actin polymerization at the barbed ends. Tension generated by myosin is transmitted along the actin bundle to the focal adhesion, where it upregulates the activity of actin polymerization module non-linearly according to Michaelis-Menten kinetics (Vavylonis et al., 2006; Jégou et al., 2013; Wu et al., 2017; Yu et al., 2017). Consistent with catch-bond behaviour of myosin observed *in vitro*, activity of myosin motors increases non-linearly as tension on the actin bundle builds up (Kovács et al., 2007). For simplicity, we ignored tension-dependent growth of focal adhesions that has been reported previously, as such growth happens on a much longer time scale (Webb et al., 2002).

When we simulated actin polymerization at a focal adhesion, we observed the emergence of an oscillating pattern of SF elongation (Fig. 5 B). According to the model, stochastic binding of myosin motors to the actin bundle leads to a gradual build-up of tension that facilitates actin polymerization at the focal adhesion driving fast SF elongation. Fast incorporation of G-actin to the barbed ends releases tension on the SF suppressing the activity of actin polymerization module and SF elongation. Thus, computational modeling of actin polymerization predicts the natural emergence of oscillatory SF elongation dynamics characterized by short periods (≈15 sec) of fast SF elongation, followed by an abrupt suppression of actin polymerization at a focal adhesion.

In addition, simulations predict that the dynamics of SF elongation is controlled by myosin-generated contractile forces. High myosin activity results in rapid tension build-up triggering a spike of actin polymerization at a focal adhesion and subsequent release of tension. Fast tension build-up results in a higher frequency of triggering events and thus, a higher average rate of SF elongation (Fig. 5 C). In contrast, low myosin activity prolongs slow SF elongation until the desired tension is reached to trigger actin polymerization at the focal adhesion and initiate a spike of SF elongation. As myosin activity is downregulated, modeling predicts a gradual decrease in the frequency and amplitude of SF elongation spikes that results in a lower average rate of SF elongation (Fig. 5 C). Thus, the dynamics of actin polymerization at a focal adhesion is predicted to exhibit myosin-dependent fluctuations that result in faster SF elongation with more frequent spikes in actin polymerization rate at a high myosin activity and slow SF elongation with rare spikes of actin polymerization when myosin contractility is supressed.

To test these model predictions, we first measured the instantaneous velocity of SF elongation. For that, we visualized retrograde movement of the photo-marker along the SF and used a Gaussian fit to locate the centroid position of the photo-marker with an accuracy of 2.3 ± 0.8 nm. Using spinning disk confocal imaging in combination with a computational sub-pixel tracking algorithm, we visualized SF elongation to derive actin polymerization dynamics at focal adhesions at 0.6 sec and 1.5 sec frame rates and found no difference in the instantaneous velocity of SF elongation (Fig. S5 B).

When we examined the detailed traces of the photo-marker, we found that SF elongation is a non-steady process with short periods of fast movement of the fluorescent photo-marker intermitted with periods of its slow movement, indicating fluctuating dynamics of actin polymerization at focal adhesions (Fig. 5 D). Using variance analysis, we assessed the variability of the photo-marks movement. For control cells the variance increased rapidly within a 5 sec window and then reached a plateau, indicating temporal fluctuations in SF elongation rate on an ≈10 sec timescale (Fig 5, E - G). Quantitative analysis revealed robust fluctuations in the instantaneous SF elongation velocity between ≈3.5 and ≈8.8 nm/sec (Fig. 5 D’). The average period of the fluctuations was 9.7 ± 3.2 sec, with most cycles of SF elongation been completed in 7 – 11 sec (Fig. 5 J). The magnitude of fluctuations in SF elongation rate was significantly larger than the experimental noise measured by tracking the fluorescent photo-markers on SFs of chemically fixed cells (Fig. S5 B) or stochastic fluctuations of SF elongation in jasplakinolide treated cells (not shown). The maximum instantaneous velocity of SF elongation was found to be strongly correlated with the average rate of SF elongation, supporting the model prediction that the magnitude of spikes in actin polymerization at focal adhesions determines the rate of SF elongation (Fig. S5 A).

We next sought to determine whether a suppression of mDia1-mediated actin polymerization affects the dynamics of SF elongation. We depleted mDia1 in cells co-expressing mEmerald-β-actin and mApple-paxillin, visualized movement of the fluorescent photo-marker along SFs and quantified the movement with a nanometer accuracy. Depletion of mDia1 suppressed fluctuations of actin polymerization as indicated by a significantly slower raise of variance of SF elongation velocity (Fig. 5 E). Cells depleted for mDia1 exhibited ≈50% decrease in the average maximum velocity of SF elongation, which made fluctuations of actin polymerization in ≈65% of focal adhesions statistically indistinguishable from experimental noise (Figs. 5 I and S5 C). In contrast, over 80% of focal adhesions in control cells exhibited statistically significant fluctuations of actin polymerization (Fig. S5 C). Surprisingly, cells depleted for mDia1 demonstrated a decreased coupling between the average SF elongation rate and the maximum instantaneous velocity of SF elongation, indicating that depletion of mDia1 may also affect the frequency of actin polymerization spikes (Fig. 5 A). In fact, power spectrum analysis revealed a 50% decrease in the frequency of actin polymerization spikes compared to the controls (not shown). Both the magnitude and the period of the fluctuations were completely regained when siRNA resistant human Dia1 was expressed in mDia1 depleted MEFs (Fig. 5, H - J). Together, these data reveal that mDia1-mediated actin polymerization at focal adhesions drives cycles of SF elongation.

Mechanical force is a potent regulator of mDia1 activity (Fig. 5C) (Jégou et al., 2013; Yu et al., 2017). To determine whether forces generated by the cytoskeleton regulate the cycles of actin polymerization at focal adhesions we tracked SF elongation on a nanometer scale in the control and mDia1 depleted cell treated with 0 – 10 μM nitro-blebbistatin. We found that attenuation of myosin activity with nitro-blebbistatin caused a dose-dependent decrease in the average maximum velocity of SF elongation in control but not mDia1 KD cells, supporting the notion that contractile forces upregulate actin polymerization activity of mDia1 (Fig. S5, D and D’). Furthermore, this treatment decreased the frequency of actin polymerization spikes and prolonged the time between sequential cycles of actin polymerization, suggesting that a decrease in myosin contractility increases the time needed to build up tension to activate mDia1 and initiate a spike in actin polymerization (Fig. 5 K). Collectively, these data support the prediction of the mathematical model that contractile forces trigger mDia1 activity at focal adhesions and initiate cycles of actin polymerization that drive SF elongation.

### mDia1-mediated SF elongation dampens mechanical tension on the SFs

According to the mathematical model, force-triggered cycles of actin polymerization release mechanical tension on SFs (Fig. 5, A - C). Thus, we sought to determine whether a suppression of actin polymerization at focal adhesions affects SF biomechanical properties. To test this, we implemented femtosecond laser nano-surgery to physically sever individual SFs and visualized the SF retraction in control and mDia1 KD cells expressing mEmerald-β-actin. Consistent with previous reports (Kumar et al., 2006; Kassianidou et al., 2017), a laser pulse applied to the central region of a SF produced an incision followed by a fast retraction of the severed SF ends reflecting a release of isometric tension (Fig. 6 A and Video S4). Quantitative analysis of the retraction dynamics revealed that the retraction distance, which reflects the elastic energy stored within the SF, was statistically different between control and mDia1 KD cells. While the retraction distance for the controls varied from 1.6 μm to 5.7 μm with the average value of 3.6 ± 1.2 μm, the SF retraction in mDia1 KDs was ≈50% larger (3.2 μm to 8.3 μm with the average value of 5.6 ± 1.7 μm), indicating a higher prestress in SFs of mDia1 KD cells (Fig. 6, B and C). The initial retraction rate (τ) of the severed SFs, along with the initial incision size (D_0_) and curve fitting (R^2^) was not significantly different between these conditions indicating that depletion of mDia1 has a negligible effect on the viscoelastic properties of the actin cytoskeleton (Fig. S6, A - C). Furthermore, by using naturally occurring fiduciary markers, we were able to rule out the possibility that retraction of the ablated SFs was a result of actin depolymerization (Fig. S6 D). These results, when combined with our finding that suppression of formins with SMIFH2 does not alter the actomyosin contractility, demonstrate that pulses of Dia1-mediated actin polymerization at focal adhesions dampen the mechanical tension on SFs. Such mechanism may act as a safety valve restricting mechanical tension on the actin cytoskeleton.

**Figure 6.**
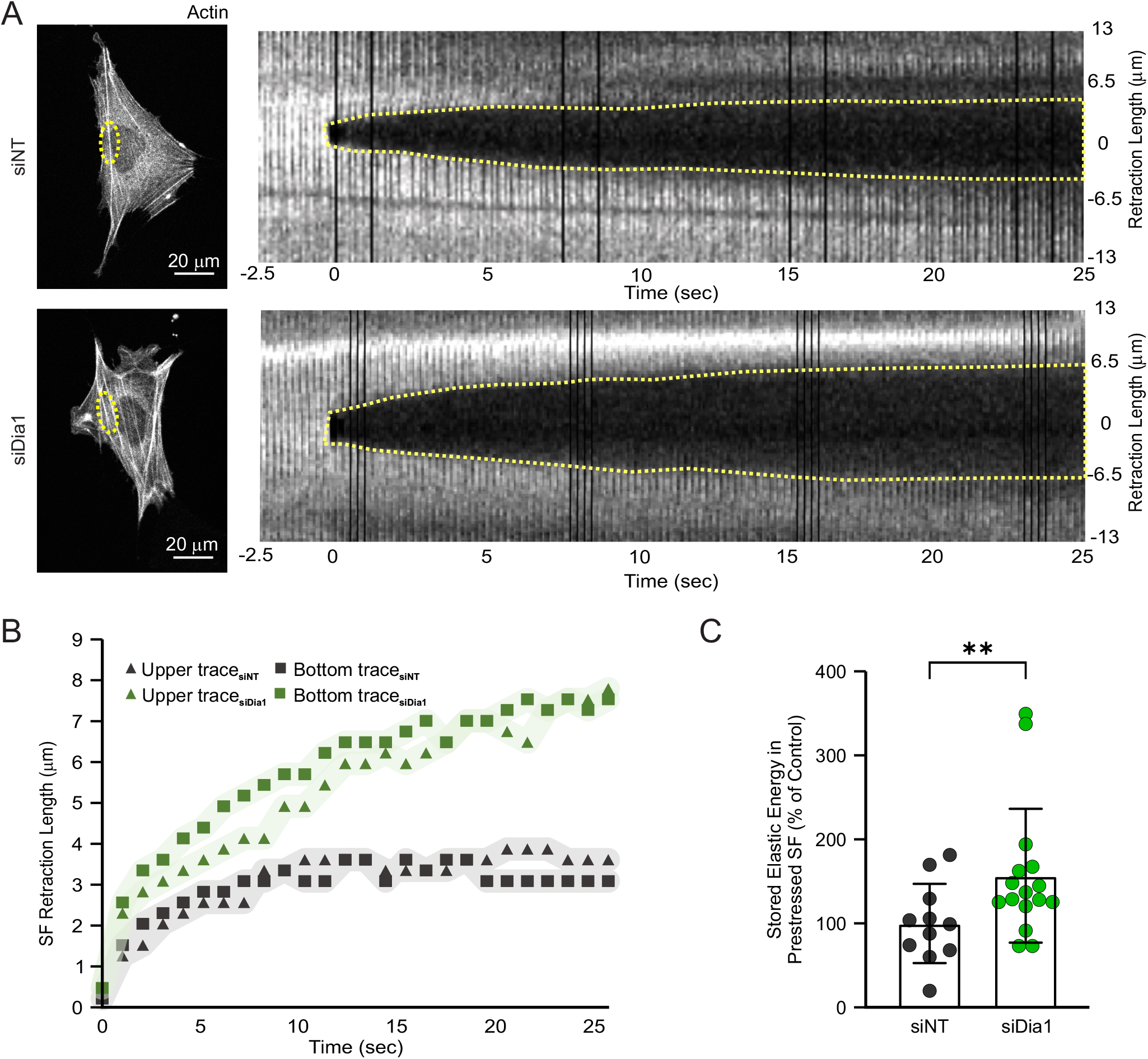
Stress fiber elongation mediated by mDia1 dampens excessive mechanical tension on the actin cytoskeleton. MEFs were transfected with non-targeting siRNA (siNT) or mDia1 targeting siRNA (siDia1) for 48 hrs and re-transfected with mEmerald-β-actin expression construct 24 hrs prior to the experiment. **(A)** Representative images of siNT and Dia1 depleted cells expressing mEmerald-actin (left panels). Corresponding montages of stress fiber images upon laser ablation (right panels). The ablated area is indicated by yellow dashed lines. **(B)** Displacement of the two newly formed stress fiber ends after laser ablation for siNT and Dia1 depleted cells. **(C)** Quantification of stored elastic energy in the prestressed stress fibers of siNT and Dia1 depleted cells. The data are presented as mean ± SD. **, p-value < 0.01; n.s., p-value > 0.05. n > 10 cells per experimental condition.

### SF elongation by mDia1 protects SFs from mechanical damage

Mechanical tension is a potent regulator of cytoskeleton organization, but excessive tension is shown to induce SF damage (Smith et al., 2010). Thus, we hypothesized that mDia1-mediated tension dampening may protect the cytoskeleton from mechanical damage and/or facilitate the efficiency of cytoskeleton repair. To test this hypothesis, we first assessed how suppression of mDia1-mediated actin polymerization affects the frequency of spontaneous cytoskeleton damage. To this end, we used time-lapse confocal microscopy to visualize the recruitment of zyxin-eGFP to the SF strain sites in mApple-actin expressing MEFs transfected with either non-targeting or mDia1-targeting siRNAs. As reported previously (Smith et al., 2010; Oakes et al., 2017), zyxin-eGFP localized to the focal adhesions and periodic puncta along SFs and such localization was not affected by mDia1 depletion (Fig. 7, A and B).

**Figure 7.**
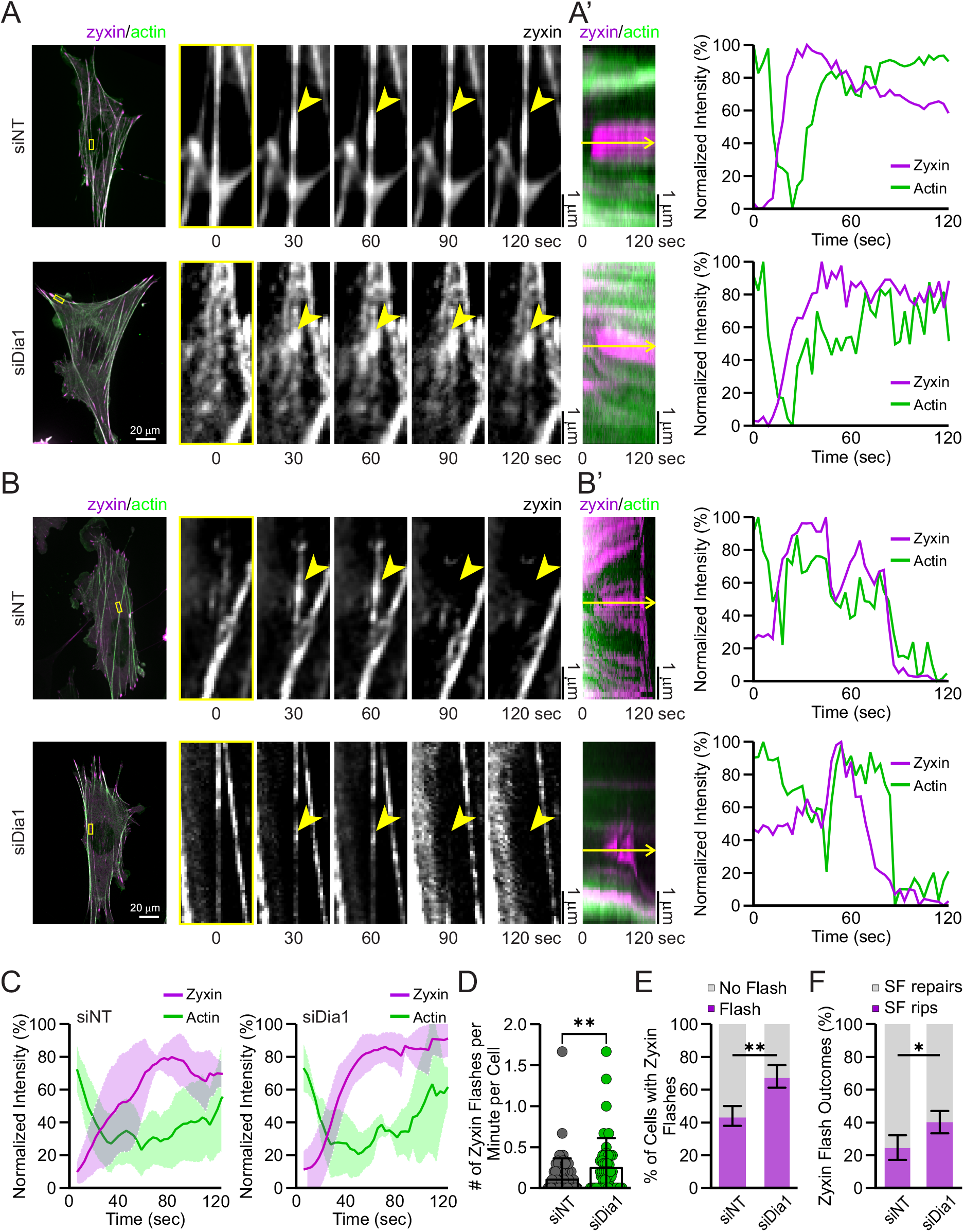
Stress fiber elongation mediated by mDia1 protects the actin cytoskeleton from mechanical damage. MEFs were transfected with non-targeting siRNA (siNT) or mDia1 targeting siRNA (siDia1) for 48 hrs and re-transfected with eGFP-zyxin and mCherry-β-actin expression constructs 24 hrs prior to the experiment. **(A)** Representative images of control and Dia1 depleted cells (left) and magnified images of the stress fibers (right) that underwent successful zyxin-mediated repair. Magnified images show the region of interests highlighted with the yellow boxes. Regions of the SFs with zyxin accumulation are indicated by the yellow arrowheads. **(A’)** Corresponding kymographs (left) and fluorescent intensity graphs (right) of highlighted regions of interest in (A). **(B)** Representative images of control and Dia1 depleted cells (left) and magnified images of the stress fibers (right) that underwent un-successful zyxin-mediated repair (right). Magnified images show the region of interests highlighted with the yellow boxes. Regions of the SFs with zyxin accumulation are indicated by the yellow arrowheads. **(B’)** Corresponding kymographs (left) and fluorescent intensity graphs (right) of highlighted regions of interest in (B). **(C)** Plot of eGFP-zyxin and mCherry-intensity over time in control (siNT, left) and Dia1 depleted cells (siDia1, right). **(D)** Bar graph of zyxin flash frequency in control and mDia1 depleted cells (n > 50 cells per experimental condition). **(E)** Stacked bar graph of the percentage of control and mDia1 depleted cells exhibiting zyxin flashes. **(F)** Quantification zyxin flash outcomes as a successful stress fiber repair or permanent damage for control and mDia1 depleted cells. n > 20 cells. The data are presented as mean ± SD. **, p-value < 0. 01; *, p-value < 0.05. n > 10 cells per experimental condition.

Additionally, zyxin-eGFP accumulated at SF strain sites identified by an acute, local decrease in actin intensity followed by an increase in zyxin-eGFP intensity (Fig. 7 A). Kymograph analysis revealed similar kinetics of zyxin recruitment to the SF strain sites in control and mDia1 KD cells (Fig. 7 C). Nevertheless, depletion of mDia1 increased SF damage, as indicated by ≈50% increase in the number of cells exhibiting zyxin recruitment to the SF strain sites and by an increase in frequency of zyxin flashes (Fig. 7 D and E). These data suggest that cells depleted of mDia1 are more susceptible to spontaneous SF damage.

To further characterize how mDia1-mediated tension dampening protects the actin cytoskeleton from mechanical damage, we determined the outcome of zyxin recruitment to the SF strain sites as either a successful repair or permeant SF damage. In some instances, recruitment of zyxin-eGFP to the SF strain sites did not lead to SF repair but followed by a catastrophic break of the SF (Fig. 7 B). Quantitative analysis of the breakage events revealed that of the SFs exhibiting zyxin accumulation at the strain sites, 75 ± 7% repaired successfully and 25 ± 7% underwent a catastrophic break in control cells. The efficiency of SF repair was significantly lower in mDia1 depleted cells with 41 ± 6% of zyxin positive strain sites underwent permanent damage and 59 ± 6% repaired successfully (Fig. 7 F). Taken together, these results demonstrate that mDia1-mediated tension dampening protects SFs from mechanical damage and facilitates cytoskeleton repair.

## DISCUSSION

Our data reveal a previously unrecognized mechanism of cytoskeleton protection whereby tension-controlled actin polymerization at focal adhesions acts as a safety valve dampening excessive mechanical tension on the actin cytoskeleton and safeguarding SFs against mechanical damage. By combining live-cell imaging with pharmacological and genetic manipulations, we identify mDia1, a member of the formin family, to be the major factor facilitating actin polymerization at focal adhesions. We demonstrate that suppression of myosin contractility results in a dose-dependent decrease in actin polymerization rate in control but not in Dia1-depleted cells, suggesting that contractility modulates mDia1 activity. Furthermore, we show that actin polymerization at focal adhesions exhibits pulsatile dynamics where the spikes of mDia1 activity are triggered by mechanical force. We demonstrate that suppression of actin polymerization at focal adhesions increases mechanical tension on SFs leading to a higher rate of spontaneous SF damage and less efficient SF repair.

Previous reports have demonstrated dynamic actin polymerization at focal adhesions and identified several actin elongating factors, including Arp2/3, formins, and Ena/Vasp family members, that might be responsible for this process (Hotulainen and Lappalainen, 2006; Iskratsch et al., 2013; Chorev et al., 2014; Puleo et al., 2019). Surprisingly, we found that formin mDia1 is the major actin elongating factor promoting actin polymerization at focal adhesions. Suppression of mDia1 activity with either a pan-formin inhibitor SMIFH2 or by depleting the protein with siRNA results in nearly complete suppression of SF elongation. A similar effect of Dia1 depletion on the rate of dorsal SF elongation has been previously reported for human osteosarcoma cells (Hotulainen and Lappalainen, 2006), implying the key role of Dia1 in actin dynamics at focal adhesions among various cell types.

Our results suggest that mechanical force is a critical regulator of mDia1 at focal adhesions. Attenuation of myosin contractility causes a dose-dependent decrease in the average rate of actin polymerization at focal adhesions in control but not mDia1-depleted cells, suggesting that contractile forces generated by the actomyosin cytoskeleton facilitate the activity of mDia1. Our results are consistent with previous theoretical and single-molecule studies showing a significant increase in the rate of mDia1-mediated actin polymerization in the presence of a physiological range of mechanical forces (Kozlov and Bershadsky, 2004; Jégou et al., 2013; Yu et al., 2017). Furthermore, we found that mechanical interactions between mDia1-mediated actin polymerization machinery and force-generating cytoskeleton predicted by mathematical modeling (Wu et al., 2017) play an important role in the regulation of mDia1 activity enabling cytoskeleton tension to serve as a gatekeeper for actin polymerization at focal adhesions.

So, what is the role of mDia1 activation by mechanical force? We propose that mDia1 acts as a safety valve releasing excessive tension on individual SFs and thus protecting the actin cytoskeleton from mechanical damage. Previous studies have shown that reduction of Dia1 expression leads to a suppression of SF assembly and a drastic reduction in the number of prominent SFs per cell (Hotulainen and Lappalainen, 2006). Despite the strong effect on the organization of actin cytoskeleton, depletion of Dia1 causes rather moderate decrease in the phosphorylation state of myosin light chain (Oakes et al., 2012). Thus, contractile forces generated by the cytoskeleton of Dia1 depleted cells are transmitted through a limited number of SFs resulting in an elevated tension on individual SFs. In fact, our measurements of SF recoil rate upon laser ablation support this notion. We found that depletion of mDia1 has a negligible effect on the viscoelastic properties of the cytoskeleton, while significantly increases tension on the SFs. Considering previous studies showing that upregulation of cytoskeletal tension initiates adaptive responses in SF architecture (Hoffman et al., 2012), our results suggest that depletion of mDia1 might supress the cell’s ability to reinforce the actin cytoskeleton upon mechanical stimulation. Based on recent evidence showing acute SF breakages induced by either an external force applied onto a cell or by internally generated forces (Smith et al., 2010; Sun et al., 2020), we hypothesized that the elevation in cytoskeletal tension observed in mDia1 depleted cells are likely to facilitate spontaneous SF damage. Surprisingly, we found that suppression of mDia1 activity does not only increase the rate of SF damage, but also decreases the efficiency of zyxin-mediated SF repair. The exact mechanisms of how depletion of mDia1 affects SF repair remains to be determined. Previous studies have demonstrated that mDia1 cooperates with Rho-dependent kinase ROCK to transform actin bundles into contractile SFs (Watanabe et al., 1999). Thus, depletion of mDia1 can result in alterations in F-actin organization in the SF, which affects the recruitment of zyxin, VASP or α-actinin to the SF strain sites (Smith et al., 2010). However, we cannot exclude that the effect of mDia1 depletion on SF breakage is purely mechanical: higher tension on the SFs leading to a faster actin thinning at the strain site that cannot be repaired by zyxin in a timely manner. Nevertheless, the interplay between mDia1 activity and zyxin-mediated cytoskeleton protection is supported by recent findings, showing that depletion of either mDia1 or zyxin perturbs the integrity of adherence junctions (Mise et al., 2012; Rao and Zaidel-Bar, 2016; Acharya et al., 2017; Harmon et al., 2021). Suppression of SF homeostasis leading to a high rate of damage might also be the primary reason for the reduced number of SFs in mDia1 depleted cells, indicating the key role of mDia1-mediated safety valve in the control of cytoskeletal architecture.

## MATERIALS AND METHODS

### Cell culture, transfection, and reagents

Mouse embryo fibroblasts (MEFs) were obtained from Dr. Beckerle (University of Utah) and primary human gingival fibroblasts (HGF) were a gift from Dr. McCulloch University of Toronto). Both MEF and HGF cells were maintained in DMEM supplemented with 2 mM L-glutamine, 100 U/mL penicillin/streptomycin and 20% FBS (Wisent Bioproducts) at 5% CO_2_. U-2 OS human osteosarcoma cells were a gift from Dr. Waterman (National Heart, Lung, and Blood Institute) and OVCAR-3 human adenocarcinoma cells were obtained from Dr. Brown (Lunenfeld-Tanenbaum Research Institute). U2 OS were cultured in McCoy’s 5A media supplemented with 2 mM L-glutamine, 100 U/mL penicillin/streptomycin and 10% FBS. OVCAR-3 cells were cultured in RPMI-1640 media supplemented with 2 mM L-glutamine, 100 U/mL penicillin/streptomycin and 20% FBS. For experiments, cells were plated on glass-bottom imaging dishes (MatTek) that had been coated with 10 μg/ml fibronectin (EMD Millipore) in PBS for 2 h at 37°C. Cells were plated at a low confluency (30–40%) in media will all the supplements and 1% FBS 16 hr prior the experiment. For experiments that require siRNA depletion or protein overexpression, cells were transfected using Nucleofector 2b (Lonza Bioscience) with a house made transfection solution (5 mM KCl, 15 mM MgCl_2_, 120 mM Na_2_HPO_4_ pH 7.2, 50 mM mannitol) and program T-020. U2 OS and OVCAR-3 cells were transfected using X-001 and T-016 programs, respectively. The following pharmacological inhibitors were used: 2.5, 5, 7.5 and 10 μM (S)-nitro-blebbistatin (Toronto Research Chemicals), 2 μM Y-27632 ROCK inhibitor (EMD Millipore), 10 μM pan-formin inhibitor SMIFH2 (EMD Millipore), 100 μM Arp2/3 inhibitor CK-666 (EMD Millipore), and 10 nM jaspakilonide (Sigma-Aldrich). The following antibodies were used: rabbit anti-paxillin, rabbit anti-phospho-myosin light chain 2 (Thr18/Ser19), rabbit anti-alpha-actinin (Cell Signalling); rabbit anti-Dia1 (Proteintech); and mouse anti-tubulin (Invitrogen). The following expression constructs were a kind gift from Michael Davidson: eGFP-actin-7 (Addgene #56421), mEmerald-actin-N-10 (Addgene #53979), mEmerald-actin-C-18 (Addgene #53978), and mApple-paxillin-22 (Addgene #54935). The expression construct for eGFP-zyxin was a gift from Dr. Waterman (National Heart, Lung, and Blood Institute). The coding sequence for human Dia1 was purchased from GeneScript and cloned in eBFP2-N1 backbone vector (gift from Michael Davidson, Addgene #54595) using XhoI/Kpn1 double digestion. Sequence of all expression constructs was confirmed by Sanger sequencing.

### Western blot analysis

Whole cell extracts of MEFs were prepared by lysing the cells in the buffer containing 50 mM Tris-HCl pH 8.0, 150 mM NaCl, 5 mM EDTA, 5% glycerol, 1% Triton X-100 supplemented with cOmplete protease inhibitor cocktail (Roche) and phosphatase inhibitor cocktails I and II (Sigma-Aldrich). Cells in the lysis buffer were mechanically ruptured by vigorous pipetting and passing through a small gauge (29G) syringe needle and the lysates were clarified by centrifugation at 24,000 *g* at 4°C for 30 minutes. Total protein concentration in the lysates was measured by BCA protein assay (Thermo). The aliquots of cell lysates containing 10 μg of total protein were mixed with a 1:4 volume of 5× Laemmli sample buffer and separated by SDS-PAGE. The proteins were transferred to an immobilon-P membrane (EMD Millipore). For protein detection, membranes were blocked for 1 hr with 2% BSA in TBS-T buffer (20 mM Tris-HCl pH 7.6, 137 mM NaCl, and 0.1% Tween-20) and incubated with primary antibodies at 4°C overnight. The membranes were washed with TBS-T and incubated with HRP-conjugated secondary antibodies for 1 hr. Protein bands were exposed to SuperSignal West Pico Chemiluminescence substrate (ThermoFisher) and visualized by an ChemiDoc detection system (BioRad). Intensity of protein bands was quantified by using ImageJ after local background subtraction.

### Collagen Contraction Assay

Collagen plug contraction assay was conducted similar to (Attieh et al., 2017). In short, MEFs (1.5 × 10^5^ cells/sample) were suspended in 2 mL of 1 mg/mL rat tail collagen I (Corning) and added to a 24-well plate in quadruplets (500 μL/well). After 1 hr of incubation at 37°C, polymerized collagen plugs were detached from the wall of the wells with a scalpel, and DMEM supplemented with 10% FBS and DMSO or nitro-blebbistatin was added. Cells were cultured for 24 hr at 37°C, 5% CO2 and images of the collagen plugs were acquired using a stereo microscope (Leica Microsystems). To quantify the collagen plug contraction, the area of the plugs was measured using ImageJ and the pe contraction was calculated as follows: 100*(Gel Area_0uM nitro-blebbistatin_/Gel Area_2.5; 5;7.5;10 μM nitro-blebbistatin_).

### Immunofluorescence

Coverslips with cells on them were fixed in 4% paraformaldehyde (Electron Microscopy Science) in PBS for 10 min and washed with TBS-T. After fixation, cells were permeabilized with 0.1 % Triton X-100 in TBS-T for 30 sec, washed and blocked in 2% BSA (BioShop) in TBS-T. Coverslips were incubated overnight at 4°C with primary antibodies diluted in the blocking solution, washed with TBS-T and incubated for 1 hr with fluorophore-conjugated secondary antibodies (Jackson ImmunoResearch Laboratories) and Alexa Fluor 488 phalloidin diluted 1:100 in the blocking solution. The stained samples were washed with TBS-T, mounted on microscope slides in aqueous mounting media (Fluoromount, Sigma-Aldrich) and sealed with a nail polish.

### Microscopy and image analysis

Images of live and immunolabeled cells were acquired on an Eclipse Ti2-E inverted microscope (Nikon Instruments) equipped with a CFI Apo TIRF 60x objective, a dynamic focusing system to correct for focus drift (PFS; Nikon Instruments), a CSU-X1 confocal scanner (Yokogawa Corporation), iChrome MLE laser combiner (Toptica Photonics), a CoolSnap Myo CCD camera (Roper Scientific), and a dual galvanometer laser scanner (Bruker Corporation). Time-lapse imaging of live cells was performed in a phenol-free DMEM media supplemented with 25 mM HEPES and 20% FBS at 37°C and at 85% humidity with a heated stage-top incubator and an objective heater (Tokai Hit).

To visualize the dynamics of actin polymerization at focal adhesions, individual SFs in mEmerald-actin and mApple-paxillin expressing cells were labeled by bleaching two parallel lines across the SF. The spacing between the lines (1 μm) was determined empirically to create a bright mEmerald spot with a Gaussian intensity profile allowing us to locate the spot with 2.4 ± 0.5 nm accuracy (Wu *et al.*, 2017). Time-lapse image series of mApple-paxillin and mEmerald-actin were acquired for 2 min at 1.5 sec or 0.6 sec frame rate and the dynamics of SF elongation was extracted from *x-t* kymographs as described (Wu *et al.*, 2017). Briefly, a kymograph of photo-marker movement was generated in NIS-Elements software (Nikon Instruments) by drawing a line along a SF starting at the center of FA anchoring the SF and averaging the mEmerald fluorescence over 0.4 μm line width. The position of a photo-label on a kymograph was tracked with sub pixel accuracy by using the Gaussian fitting function of Matlab Statistics toolbox. The confidence intervals of the tracked positions are obtained by the regression parameter confidence interval estimator in Matlab Statistics toolbox.

Area of individual focal adhesions was measured from the immunofluorescent images of cells stained with anti-paxillin antibody using a publicly available Focal Adhesion Analysis Server (Goffin et al., 2006; Gardel et al., 2010; Berginski and Gomez, 2013). Quantification of focal adhesion maturation was performed based on the area of individual focal adhesions as described (Goffin et al., 2006; Gardel et al., 2010). Small paxillin-positive adhesions (area ranges from 0.05 to 0.20 μm^2^) were classified as nascent adhesions. Focal adhesions with an area ranging from 0.20 to 1.75 μm^2^ and > 1.75 μm^2^ were classifies as mature and fibrillar adhesions, respectively. The percentage of each type of the adhesions was calculated for individual cells and then averaged for every experimental condition.

Quantitative analysis of SF morphology was performed on the images cells stained with fluorescently labelled phalloidin using an FSegement Matlab script developed by Rogge *et al.* (Rogge et al., 2017). In short, SFs were segmented by using a trace algorithm optimized to detect linear structures on the images with subtracted background fluorescence. Segmented linear structures were analyzed with respect to fluorescent values for thick and thin actin fibers as well as the total length and number of SFs.

Quantitative analysis of the cytoskeleton damage was performed as described (Smith et al., 2014). Time-lapse image series of MEFs co-transfected with eGFP-zyxin and mApple-actin expression constructs were acquired every 3 sec for 10 min. The images were inspected visually to identify and count the events of zyxin recruitment to the spontaneously damaged SFs. Zyxin frequency and outcomes where recorded for every cell and average for each experimental condition. Kinetics of zyxin recruitment to the SF strain sites was determined by kymograph analysis using NIS-Elements software (Nikon Instruments).

### Nano-surgery of single SFs

MEFs expressing mEmerald-β-actin were plated on the coverslips coated with 10 μg/mL human plasma fibronectin (EMD Millipore). The coverslips with cells were mounted on the microscope slides for imaging, filled up with phenol-free DMEM supplemented with 25 mM HEPES and 20% FBS, sealed with VALAP, and kept at 37°C for 1 hr to allow cells to acclimatize (Clarke and Braun, 2017) Image acquisition and SF photo-ablation were performed by an Eclipse FN1 upright microscope (Nikon Instruments) equipped with an A1R-MP confocal scanner and 25X CFI Apo NA 1.1 LWD objective. Images were acquired with a resonant scanner and 488nm laser line. Femtosecond infrared laser (Chameleon Vision II, Coherent) was used for SF photo-ablation. During the experiments, the cells were maintained at 37°C by with an ASI 400 air stream incubator (NevTek) and a heated objective coil (Tokai Hit).

SF nano-surgery was performed similar to the protocol published by Kumar *et al*. (2006). Briefly, the femtosecond laser was set at 800 nm and configured to make a linear scan at 30% of maximal power and 30 μsec/pixel dwell time across an individual SF. SF retraction was recorded at a frame rate of 130 msec for the first 30 sec and then at 1.6 sec for an additional 120 sec. SF retraction dynamics were quantified by kymograph analysis with NIS-Elements (Nikon Instruments). The resultant kymographs were fit to a Kelvin–Voigt model, *L*_*t*_=D_a_+L_o_[1−exp(−tτ)], where *L*_*t*_ is a half of the measured distance between the two severed SF ends; D_a_ is the length of SF destroyed by ablation; L_o_ is the retraction plateau distance; and *τ* is the viscoelastic constant (Kumar et al., 2016; Kassianidou et al., 2017).

### Mathematical model description

Based on the literature and our established work (Wu et al., 2017), the model describes the negative feedback between SF-mediated actomyosin contraction and the associated SF elongation. That is, actomyosin contractility generates tension upon the SF plus-end, which speeds up the SF elongation by incorporating the actin monomers. This SF elongation facilitates and hence, tunes down the associated actomyosin contraction (Fig. 5A).

In the spirit of simplicity, the model only describes three key components of this negative feedback that directly interface with our experiments. They are the SF contractility, the levels of actin nucleation factor and the profilin-bound actin monomer at the SF plus end, which are denoted as “C”, “N”, and “A” in the model. The model assumes that SF connects to focal adhesion, without the explicit depiction of the latter. Below is the mathematical formulation of our model.

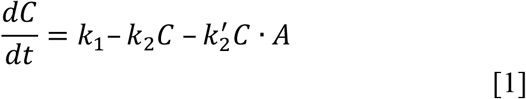

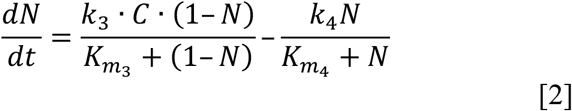

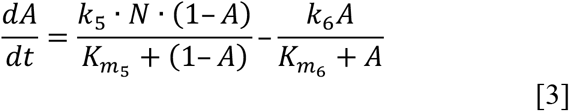

Here, Eq. [1] describes the dynamics of SF contractility with the activation rate of *k*_1_, the basal deactivation rate of *k*_2_, and importantly, the SF elongation-facilitated deactivation rate of 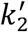. We assume that the SF elongation rate is linearly proportional to the local level of profilin-bound actin monomer (*A*), which is ready to be incorporated into SF plus end, driving the SF elongation. Eq. [2] is the Michaelis-Menten equation that depict the enzymatic activation of actin nucleation factor (*N*) by the SF contractility (*C*) and its intrinsic deactivation at a rate of *k*_4_. Likewise, Eq. [3] is the Michaelis-Menten equation describing the dynamics of the local level of profilin-bound actin monomer (*A*), which is activated by the actin nucleation factor (*N*) and its basal deactivation (the *k*_6_-term).

### Statistics

All statistical analyses were performed with GraphPad Prism 8 statistical package. For pairwise comparisons, statistical significance was calculated by Student’s t-test. For multivariance analysis we used ANOVA with Tukey’s post hoc test for pairwise comparisons of normally distributed data. Data were assumed to satisfy the ANOVA requirements if the residuals of a dataset were normally distributed (Quinn and Keough, 2002). Chi-square test was used in the comparisons of non-normally distributed variables.

As denoted on the figures, differences between the experimental groups with p-values < 0.05 were considered statistically significant. Data on the bar graphs are presented as individual measurements (data points, dots) with their mean ± standard deviation (SD).

### Online supplemental materials

Fig. S1 describes the effect of C- and N-terminally tagged actin on the SF architecture. Fig. S2 includes the quantitative analysis of fiduciary markers bleached on the SFs and a bar graph of SF density in cells treated with formin (SMIFH2) and Arp2/3 (CK-666) small molecule inhibitors. Fig. S3 illustrates the effect of mDia1 suppression on the organization and contractile properties of the actin cytoskeleton. Fig. S4 shows the effect of partial inhibition of myosin with (S)-nitro-blebbistatin on the morphology of SFs and focal adhesions. Fig. S5 illustrates the effects of mDia1 depletion and myosin inhibition on the cycles of mDia1-mediated SF elongation. Fig. S6 shows quantifications of SF viscoelastic properties in control and mDia1-depleted cells. Video 1 shows the dynamics of SF elongation at focal adhesions in control cells. Video 2 shows the dynamics of SF elongation at focal adhesions in mDia1-depleted cells. Video 3 shows the rescue of SF elongation in mDia1 depleted cells upon re-expression of siRNA resistant hDia1-BFP. Video 4 shows SF recoil after laser ablation.

## ACKNOWLEDGMENTS

We thank Tony Harris and Boris Hinz for comments on the manuscript. This work was supported by Connaught Fund New Investigator Award to S.P., Canada Foundation for Innovation, NSERC Discovery Grant Program (grants RGPIN-2015-05114 and RGPIN-2020-05881), University of Manchester and University of Toronto Joint Research Fund, and University of Toronto XSeed Program The authors declare no competing financial interests.

## AUTHOR CONTRIBUTIONS

F.R. Valencia, J. Liu, and S.V. Plotnikov conceived the study, designed and performed the experiments, and analyzed data. E. Sandoval and J. Liu developed computational analysis tools and performed data analysis. F.R. Valencia and S.V. Plotnikov wrote the manuscript with input from all authors. All authors reviewed the results and approved the final content of the manuscript.

## ABBREVIATIONS

The abbreviations used are:

SF: stress fiber
MEF: mouse embryo fibroblasts
KD: gene knockdown
BFP: blue fluorescent protein
HGF: human gingival fibroblasts

## SUPPLEMENTAL FIGURE AND VIDEO CAPTIONS

**Supplemental Figure 1.**
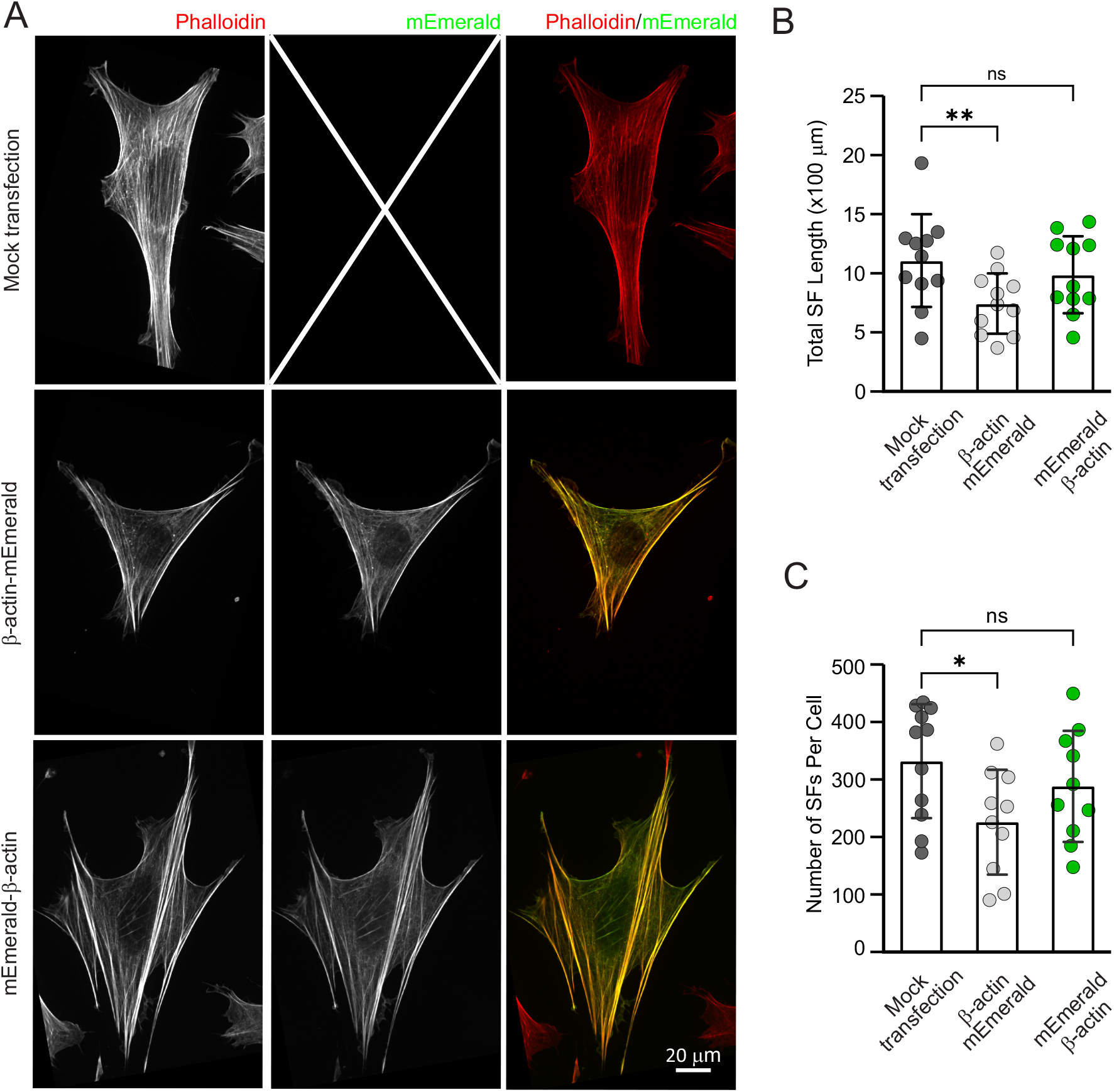
Overexpression of N-terminally tagged actin has a negligible effect on stress fiber architecture. **(A)** Representative images of mock transfected MEFs (top row) and cells expressing β-actin tagged with mEmerald at the C- or N- termini (middle and bottom row, respectively) stained with phalloidin-AlexaFluor561. **(B, C)** Bar graphs of the total stress fiber length **(B)** and the number of stress fibers **(C)** in mock transfected MEFs and cells expressing β-actin tagged with mEmerald at the C- or N-termini. Each data point in the graph represents an individual quantified cell in one of three independent experiments. The data are presented as mean ± SD. **, p-value < 0. 01; *, p-value < 0.05; n.s., p-value > 0.05. Scale bar, 20 μm. n > 11 cells per an experimental condition.

**Supplemental Figure 2.**
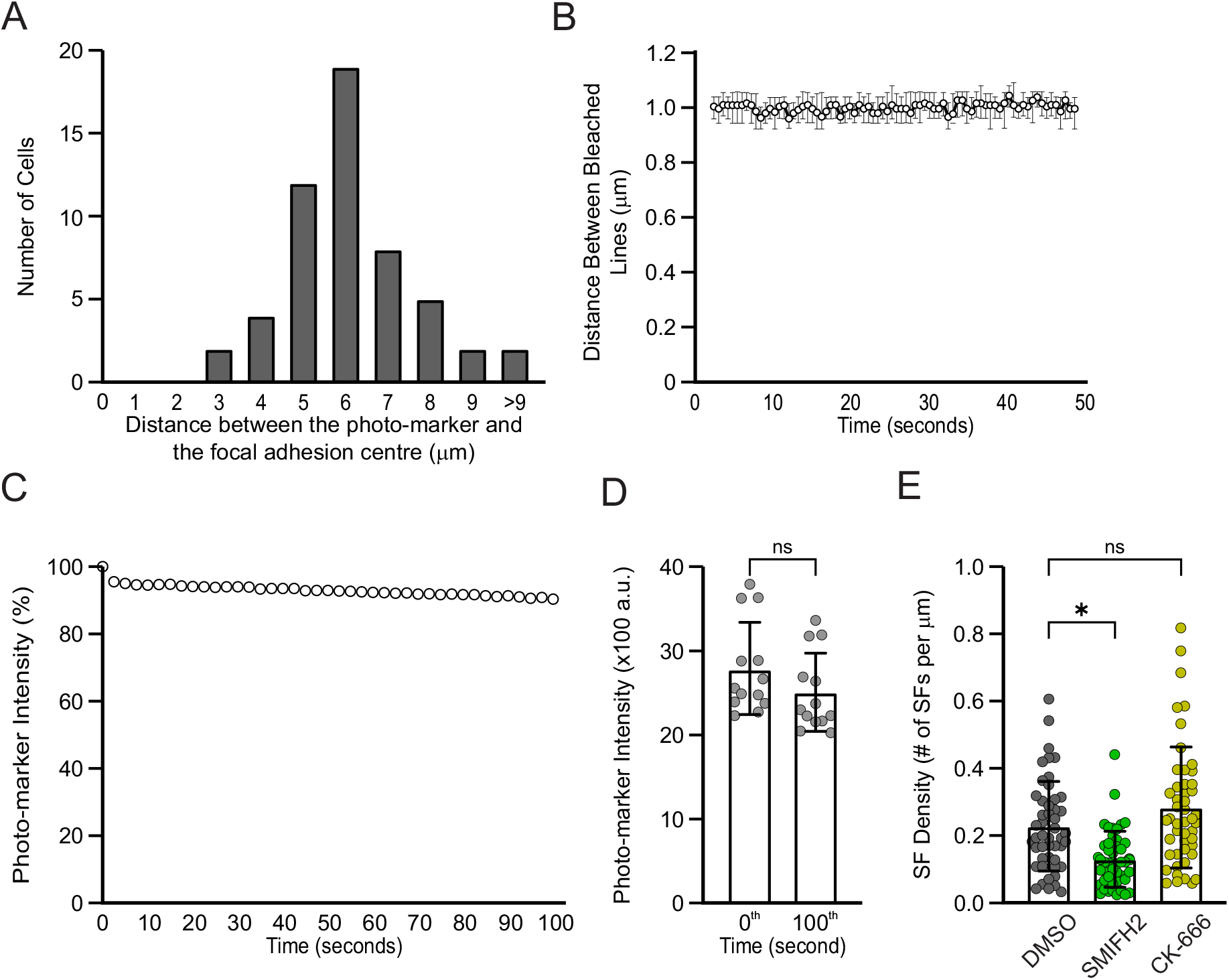
Stress fiber elongation at focal adhesions requires formin activity. **(A – D)** MEFs were co-transfected with mEmerald-β-actin and mApple-paxillin expression constructs 24 hrs prior to the experiment. The following characteristics of the fluorescent photo-marker on stress fiber were quantified: **(A)** distribution of the distance between the photo-marker and the center of focal adhesion where the stress fiber originates. **(B)** Average width of the fluorescent photo-marker over 100 sec time period. **(C)** Average intensity of the fluorescent photo-marker over 100 sec time period. The intensity values were normalized by the fluorescence intensity for the first frame of the movie. **(D)** Bar graph of the photo-markers’ intensity at the 0^th^ or 100^th^ second**. (E)** Bar graph of stress fiber density in control, SMIFH2- and CK-666-treated cells. Cells plated on the fibronectin-coated coverslips were treated with 0.1% DMSO (Control), 10 μM pan-formin inhibitor SMIFH2, 100 μM Arp2/3 inhibitor CK-666 for 1 hr, fixed and immunostained with phalloidin-AlexaFluor488. Each data point in the graph represents an individual quantified cell in one of three independent experiments. The data are presented as mean ± SD. *, p-value < 0.05; n.s., p-value > 0.05. n > 12 cells per an experimental condition.

**Supplemental Figure 3.**
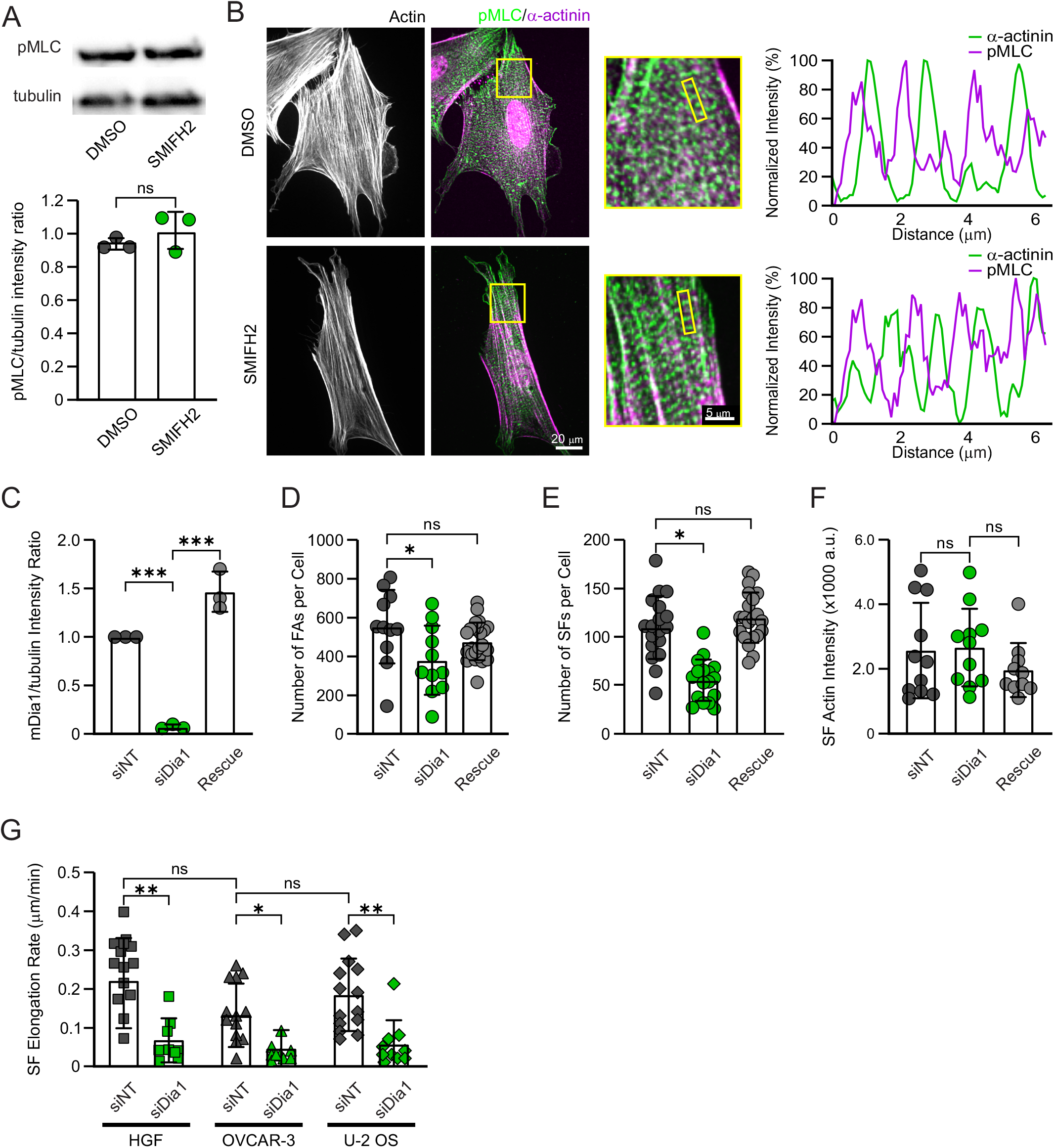
Suppression of mDia1 has a minimal effect on the organization of actomyosin cytoskeleton. **(A, B)** MEFs were treated with 0.1% DMSO (vehicle) or 10 μM pan-formin inhibitor SMIFH2 for 1 hr. **(A)** Representative Western blot (above) and corresponding bar graph (below) of total cell lysates stained with the antibodies against phosphorylated myosin light chain (pMLC) and tubulin. For the bar graph, the intensity of pMLC band was normalized by the tubulin band’s intensity. The average pMLC intensity ± SD calculated from three independent Western blots is presented. **(B)** Representative images of MEFs stained with fluorescently labeled phalloidin (left panels) and the antibodies against pMLC and α-actinin. Zoom-ins of selected regions highlighted by the yellow boxes are shown on the right. Right panels: Profiles of the fluorescence intensity along the yellow lines **(B)** demonstrates sarcomere-like pattern in control (top panel) and SMIFH2-treated (bottom panel) cells. **(C - F)** MEFs were transfected with non-targeting (Control) or mDia1-targeting (siDia1 and Rescue) siRNAs 48 hrs prior the experiments. For the rescue condition cells were additionally transfected with hDia1-BFP expression construct 24 hrs after siRNA transfection. **(C)** Quantification of Dia1 band intensity on the Western blot of total cell lysates stained with the antibodies against mDia1 and tubulin. The intensity of Dia1 band was normalized to the tubulin intensity. The average Dia1 intensity ± SD calculated from three independent Western blots is presented. **(D - F)** Bar graphs of the number of focal adhesions **(D)** and stress fibers **(E**), and the intensity of stress fibers stained with fluorescently labeled phalloidin **(F)** in control, mDia1-depleted and hDia1-rescued cells. Each data point in the graph represents an individual quantified cell in one of three independent experiments. n > 25 cells per an experimental condition. **(G)** Bar graph of the average rate of stress fiber elongation in primary human gingival fibroblasts (HGF), human ovarian adenocarcinoma (OVCAR-3), and human osteosarcoma (U-2 OS) cells. n > 8 cells per an experimental condition. The data are presented as mean ± SD. ***, p-value < 0.001; **, p-value < 0.01; *, p-value < 0.05; n.s., p-value > 0.05. Scale bar for whole cell images is 20 μm and for zoom-ins is 5 μm.

**Supplemental Figure 4.**
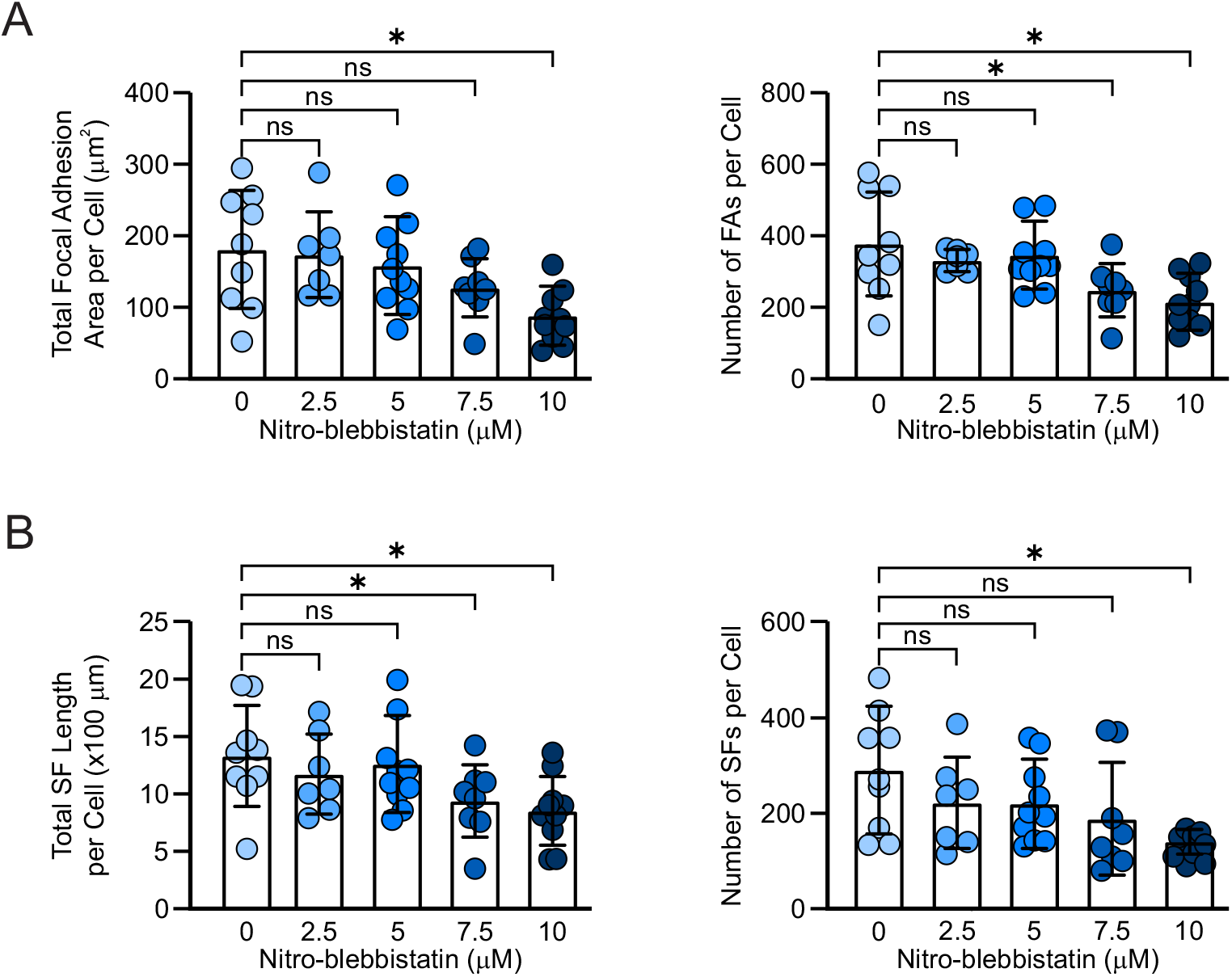
Partial inhibition of myosin contractility with (S)-nitro-blebbistatin results in a moderate suppression of stress fiber and focal adhesion assembly. **(A, B)** MEFs plated on fibronectin-coated coverslips were treated with either 0.1% DMSO (vehicle) or 2.5 μM, 5 μM, 7.5 μM, 10 μM (S)-nitro-blebbistatin for 2 hrs followed by immunostaining for filamentous actin and focal adhesions. Confocal images of the cells were analyzed quantitatively to assess changes in organization of the actin cytoskeleton and focal adhesions (see Methods for details). **(A)** Bar graphs of the total focal adhesion area per cell (left) and the number of focal adhesions per cell (right). (**B)** Bar graphs of the total stress fiber length per cell (left) and the number of stress fibers per cell (right). The data are presented as mean ± SD. *, p-value < 0.05; n.s., p-value > 0.05. n > 7 cells per an experimental condition.

**Supplemental Figure 5.**
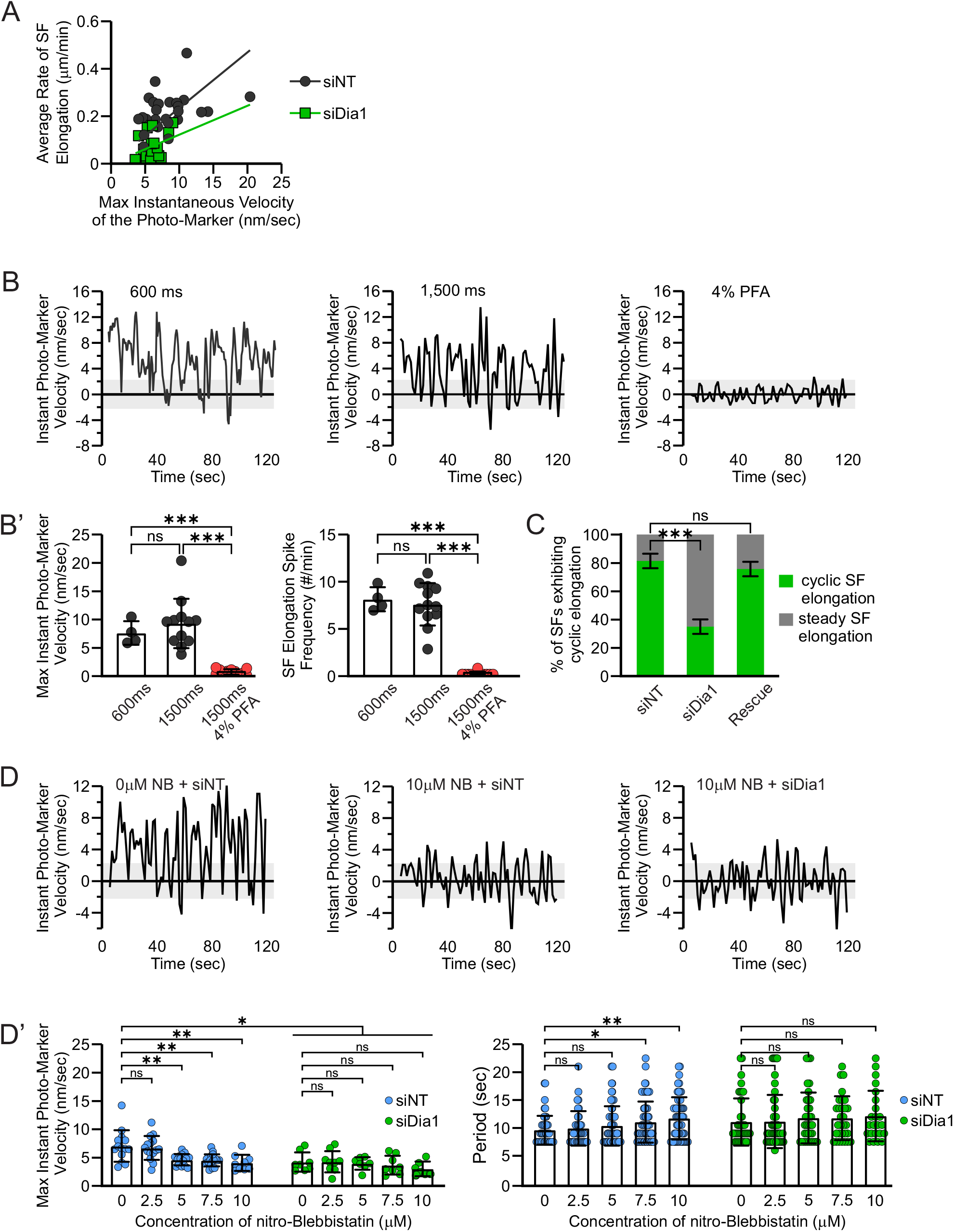
Depletion of mDia1 supresses cycles of stress fiber elongation at focal adhesions. MEFs were transfected with either non-targeting (siNT) or mDia1 (siDia1) targeting siRNA 48 hrs prior to the experiments. To visualize dynamics of stress fiber elongation, the cells were additionally co-transfected with the siRNA listed above and the expression constructs for mEmerald-β-actin and mApple-paxillin 24 hrs prior to the experiments. **(A)** Plot of average rate *vs* maximal instantaneous velocity of stress fiber elongation at individual focal adhesions in control and mDia1 depleted cells. **(B)** Representative profiles of stress fiber elongation rate for movies acquired at 600 ms (left) and 1500 ms (middle and right) frame rate. To determine the contribution of experimental noise, movement of the photo-marker was quantified in cells chemically fixed with 4% paraformaldehyde (right panel). **(B’)** Quantification of maximum instantaneous rate of stress fiber elongation (left) and frequency of stress fiber elongation spikes (right) measured for movies acquired at 600 ms and 1500 ms frame rate. **(C)** Stacked bar graph of the percentage of stress fibers exhibiting cyclic elongation in cells transfected with non-targeting siRNA (siNT) or mDia1-targeting siRNA (siDia1 and Rescue). For the rescue conditions cells were additionally transfected with siRNA resistant hDia1-BFP expression construct. **(D)** Representative profiles of stress fiber elongation rate in cells transfected with either non-targeting siRNA (siNT; left and middle) or mDia1-targetting siRNA (siDia1; right panel) treated with 0.1% DMSO (vehicle, left) or 10 μM nitro-blebbistatin (middle and right). **(D’)** Quantification of maximum instantaneous rate of stress fiber elongation (left) and period of stress fiber elongation spikes in control and mDia1 depleted cells treated with a range (0 – 10 μM) of nitro-blebbistatin. The data on the bar graphs are presented as mean ± SD. ***, p-value < 0.001; **, p-value < 0. 01; *, p-value < 0.05; n.s., p-value > 0.05. For B’, n > 5 cells per an experimental condition. For C’, n > 10 cells per an experimental condition.

**Supplemental Figure 6.**
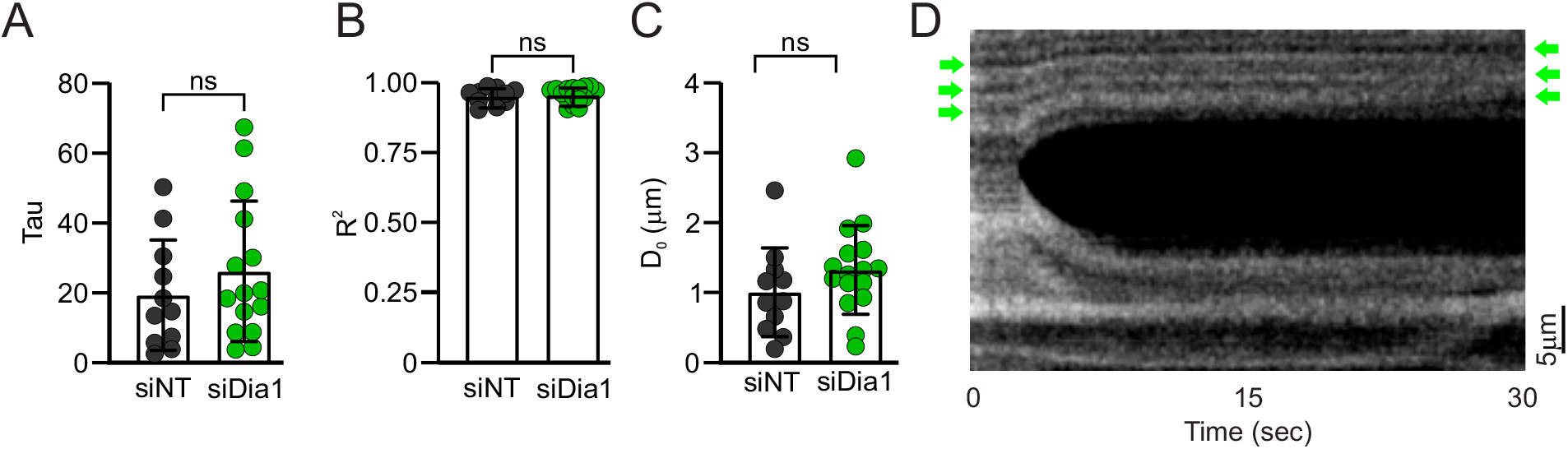
Retraction of stress fiber upon nano-surgery is due to changes in mechanical tension. Quantification of **(A)** the initial retraction rate (τ) of the ablated stress fibers, **(B)** along with the initial incision size (D_0_) and **(C)** curve fitting (R^2^) of viscoelastic properties for control and mDia1 depleted cells. n.s., p-value > 0.05. n > 15 cells per an experimental condition. **(D)** Kymograph of stress fiber undergoing nano-surgery with naturally occurring fiduciary markers indicated by green arrows at the beginning and end of the acquisition.

Supplemental Video 1. **Actin polymerization at focal adhesions of a MEFs treated with non-targeting siRNA**. Actin dynamics at focal adhesions in a MEF cell treated with non-targeting siRNA (siNT), visualized by expression of mEmerald-β-actin(green) and mApple-paxillin (red). The time lapse images were acquired every 1.5 seconds for 2 minutes. Frame rate is 10 frames per second. To the left is a frame of the whole cell – scale bar is 20 μm – and to the right is the corresponding zoomed in region denoted by a yellow box of a photo-label on a stress fiber moving away from its corresponding focal adhesion – scale bar is 2 μm. The yellow arrow indicates the first position of the photo-label in the movie and the red arrow the last position of the photo-label in the movie. Related to Fig 3, C and D).

Supplemental Video 2. **Actin polymerization at focal adhesions of a MEFs treated with mDia1-targeting siRNA**. Actin dynamics at focal adhesions in a MEF cell treated with mDia1-targeting siRNA (siDia1), visualized by expression of mEmerald-β-actin(green) and mApple-paxillin (red). The time lapse images were acquired every 1.5 seconds for 2 minutes. Frame rate is 10 frames per second. To the left is a frame of the whole cell – scale bar is 20 μm – and to the right is the corresponding zoomed in region denoted by a yellow box of a photo-label on a stress fiber moving away from its corresponding focal adhesion – scale bar is 2 μm. The yellow arrow indicates the first position of the photo-label in the movie and the red arrow the last position of the photo-label in the movie. Related to Fig 3, C and D).

Supplemental Video 3. **Actin polymerization at focal adhesions of a mDia1 depleted MEF cell rescued by re-expression of hDia1**. Actin dynamics at focal adhesions in a mDia1 depleted MEF cell rescued by re-expression of hDia1 (Rescue), visualized by expression of mEmerald-β-actin(green) and mApple-paxillin (red). The time lapse images were acquired every 1.5 seconds for 2 minutes. Frame rate is 10 frames per second. To the left is a frame of the whole cell – scale bar is 20 μm – and to the right is the corresponding zoomed in region denoted by a yellow box of a photo-label on a stress fiber moving away from its corresponding focal adhesion – scale bar is 2 μm. The yellow arrow indicates the first position of the photo-label in the movie and the red arrow the last position of the photo-label in the movie. Related to Fig 3, C and D.

Supplemental Video 4. **Stress fiber elongation mediated by mDia1 dampens excessive mechanical tension on the actin cytoskeleton.** Nano-surgery of stress fibers of MEFs transfected with non-targeting siRNA (siNT) to the left or mDia1 targeting siRNA (siDia1) for 48 hrs and re-transfected with mEmerald-β-actin expression construct 24 hrs prior to the experiment to the right. The time lapse images were acquired every 0.26 seconds for the first 30 seconds and then every 500 milliseconds for 1 minute after. Frame rate is 50 milliseconds per frame. The scare bar is 20 μm. Red eclipses indicate areas of stress fiber recoil. Related to Fig 6 C.

